# Active zone protein SYD-2/Liprin-α acts downstream of LRK-1/LRRK2 to regulate polarized trafficking of synaptic vesicle precursors through clathrin adaptor protein complexes

**DOI:** 10.1101/2023.02.26.530068

**Authors:** Sravanthi S P Nadiminti, Shirley B Dixit, Neena Ratnakaran, Sneha Hegde, Sierra Swords, Barth D Grant, Sandhya P Koushika

## Abstract

Synaptic vesicle proteins (SVps) are thought to travel in heterogeneous carriers dependent on the motor UNC-104/KIF1A. In *C. elegans* neurons, we found that some SVps are transported along with lysosomal proteins by the motor UNC-104/KIF1A. LRK-1/LRRK2 and the clathrin adaptor protein complex AP-3 are critical for the separation of lysosomal proteins from SVp transport carriers. In *lrk-1* mutants, both SVp carriers and SVp carriers containing lysosomal proteins are independent of UNC-104, suggesting that LRK-1 plays a key role in ensuring UNC-104-dependent transport of SVps. Additionally, LRK-1 likely acts upstream of the AP-3 complex and regulates the membrane localization of AP-3. The action of AP-3 is necessary for the active zone protein SYD-2/Liprin-α to facilitate the transport of SVp carriers. In the absence of the AP-3 complex, SYD-2/Liprin-α acts with UNC-104 to instead facilitate the transport of SVp carriers containing lysosomal proteins. We further show that the mistrafficking of SVps into the dendrite in *lrk-1* and *apb-3* mutants depends on SYD-2, likely by regulating the recruitment of the AP-1/UNC-101. We propose that SYD-2 acts in concert with both the AP-1 and AP-3 complexes to ensure polarized trafficking of SVps.

## Introduction

Synaptic vesicles (SVs) found at the pre-synaptic terminal contain membrane-associated proteins, such as Synaptobrevin-1 (SNB-1), Synaptogyrin-1 (SNG-1), SV2, and RAB-3 (Takamori *et al*., 2006). They are known to have a well-defined composition lacking, for instance, Golgi-resident enzymes (Salazar *et al*., 2004; Takamori *et al*., 2006; Choudhary *et al*., 2017). The loss of SV proteins (SVps) has been shown to affect neurotransmission (Nonet *et al*., 1998; Mahoney, Luo and Nonet, 2006; Brockmann *et al*., 2020; Richmond, Davis and Jorgensen, 1999; Aravamudan *et al*., 1999) and the progression of neurodegenerative disorders (Kraemer *et al*., 2003). However, the trafficking routes of SVps in the cell body remain to be fully elucidated. Although SNB-1 and SNG-1 are present along with RAB-3 at synapses, only a subset of the SNB-1 and SNG-1 carriers that exit the cell body includes RAB-3 (Choudhary *et al*., 2017; Maeder, Shen and Hoogenraad, 2014). Likewise, Synaptophysin and SV2 do not appear to be co-transported by the mammalian SV motor KIF1A (Okada and Hirokawa, 1999), while Synaptophysin and the Zinc transporter ZnT3 are likely enriched in different populations of synaptic-like microvesicles (Salazar *et al*., 2004). Additionally, SVp carriers exiting the cell body are tubular as opposed to those closer to the synapse, which have a defined smaller diameter (Tsukita and Ishikawa, 1980; Nakata, Terada and Hirokawa, 1998). Prior studies from mammalian cells and *Drosophila* suggest that some SVps share trafficking routes with lysosomal proteins (Newell-Litwa *et al*., 2009; Vukoja *et al*., 2018; Rizalar, Roosen and Haucke, 2021). These findings suggest that SVps emerge from the cell body in precursor or immature transport carriers that likely have a heterogeneous composition, sharing trafficking routes with lysosomal proteins.

Several genes have been identified as important in the trafficking of SVps. UNC-16/JIP3-mediated recruitment of LRK-1/LRRK2 on the Golgi seems to be critical for excluding Golgi-resident enzymes from SVp carriers as well as regulating the size of these carriers (Choudhary *et al*., 2017). The AP-3 complex has been shown to play a key role in separating SVps and lysosomal proteins that initially occupy a common intermediate compartment (Newell-Litwa *et al*., 2009). The biogenesis and maturation of precursor vesicles containing the endolysosomal protein LAMP-1, active zone proteins, and SV proteins are regulated by RAB-2 (Götz *et al*., 2021). UNC-104/KIF1A is the kinesin motor important for SVp transport (Hall and Hedgecock, 1991; Okada and Hirokawa, 1999; Zhao *et al*., 2001; Pack-Chung *et al*., 2007). We previously showed that the SVp carriers formed in the *unc-16*/*jip3*, *lrk-1*/*lrrk2*, and *apb-3* (mutant of the β subunit of the AP-3 complex) mutants of *Caenorhabditis elegans* are not exclusively dependent on UNC-104/KIF1A for their transport (Choudhary *et al*., 2017). However, the link between the maturation of SVp carriers and their ability to recruit the SVp motor remains to be well understood.

Active zone proteins that mark release sites for SVs at synapses have also been shown to co-transport with some SVps (Bury and Sabo, 2011; Xuan *et al*., 2017; Vukoja *et al*., 2018; Lipton, Maeder and Shen, 2018). Moreover, SVps and some active zone proteins, such as ELKS-1, have been shown to co-transport in lysosomal protein-containing packets called presynaptic lysosome-related vesicles (PLVs). These PLVs are dependent on the small GTPase ARL-8, an interactor of UNC-104/KIF1A/IMAC, which is thought to facilitate UNC-104/KIF1A interaction with the PLVs (Vukoja *et al*., 2018). Additionally, active zone proteins Piccolo and Bassoon present in clusters with Synaptobrevin, Synaptotagmin, and SV2, are thought to be important in forming such transport clusters (Tao-Cheng, 2020). Together, these data suggest that SVp and lysosomal protein trafficking and transport can be regulated by active zone proteins.

SYD-2/Liprin-α, an active zone protein, is known to interact with and bind to the SV motor UNC-104/KIF1A (Zheng *et al*., 2014; Shin *et al*., 2003; Wagner *et al*., 2009; Stucchi *et al*., 2018). SYD-2/Liprin-α also influences the distribution of acidic organelles such as SVs (Zheng *et al*., 2014), dense core vesicles (Goodwin and Juo, 2013), and lysosomes (Edwards *et al*., 2015b). Active zone proteins SYD-2 and SYD-1 along with synapse assembly proteins SAD-1 and CDK-5 are known to regulate lysosomal protein trafficking in *unc-16* mutants through dynein (Edwards *et al*., 2015b). ELKS-1, which binds SYD-2 (Ko *et al*., 2003; Dai *et al*., 2006), has been shown to interact with RAB-6 to regulate the trafficking of melanosomal proteins (Patwardhan *et al*., 2017) and SVs (Nyitrai, Wang and Kaeser, 2020). These studies suggest that SYD-2 can affect the trafficking of SVs and other acidic organelles.

In this study, we used the *C. elegans* touch receptor neuron (TRN) model to better define the co-transport and eventual separation of SV and lysosomal proteins. Importantly, we show that LRK-1 and the AP-3 complex, which we previously identified as important for regulating SV precursor composition (Choudhary *et al*., 2017), play a critical role in sorting lysosomal proteins away from SVps. Furthermore, the active zone protein SYD-2/Liprin-α plays a key role along with UNC-104/KIF1A in the transport of compartments containing both SVps and lysosomal proteins in the absence of the AP-3 complex. Our data suggest that although the SV motor can be recruited on compartments that contain both SVps and lysosomal proteins, SV precursors lacking lysosomal proteins appear to preferentially recruit the SV motor UNC-104.

## Results

### Synaptic vesicle proteins travel with lysosomal proteins in heterogenous carriers

Although studies have indicated that SVps are transported in heterogeneous carriers, the composition of these carriers has not been fully examined. Here, we assessed the co-transport of specific SVps with one another and with other endomembrane compartment proteins in the proximal posterior lateral mechanosensory (PLM) neuron of *C. elegans* (Fig. S1A, S1B-G; Movies S1 and S2).

Less than 10% of moving RAB-3- and MAN-II-containing vesicles co-transport both markers (Fig. 1A). Likewise, ∼10% of SNG-1 and SNB-1 are co-transported with the lysosome-specific cystine transporter, cystinosin (CTNS-1) (Kalatzis *et al*., 2001) (Fig. 1A and Movie S1). Nearly all CTNS-1- and RAB-7-carrying compartments co-transport SNG-1, while only ∼50% of CTNS-1-labelled compartments co-transport SNB-1 (Fig. 1B). CTNS-1-labelled compartments move in both the anterograde and retrograde directions in wildtype (Fig. 1C). Approximately 30% of SNG-1 and RAB-7 are co-transported (Fig. 1A, Movie S2). Furthermore, nearly every CTNS-1-carrying compartment co-transports RAB-7, while only ∼40% of RAB-7-carrying compartments co-transport CTNS-1 (Fig. 1D). RAB-3, a synaptic vesicle RAB, does not co-transport with CTNS-1, while RAB-3 and RAB-7 are co-transported ∼10% of the time (Fig. 1A). RAB-3 is co-transported with SNB-1 and SNG-1 approximately 35% of the time (Fig. 1A). Thus, SVp carriers exiting the cell body largely exclude the Golgi-resident enzyme MAN-II and lysosomal proteins.

**Figure 1:**
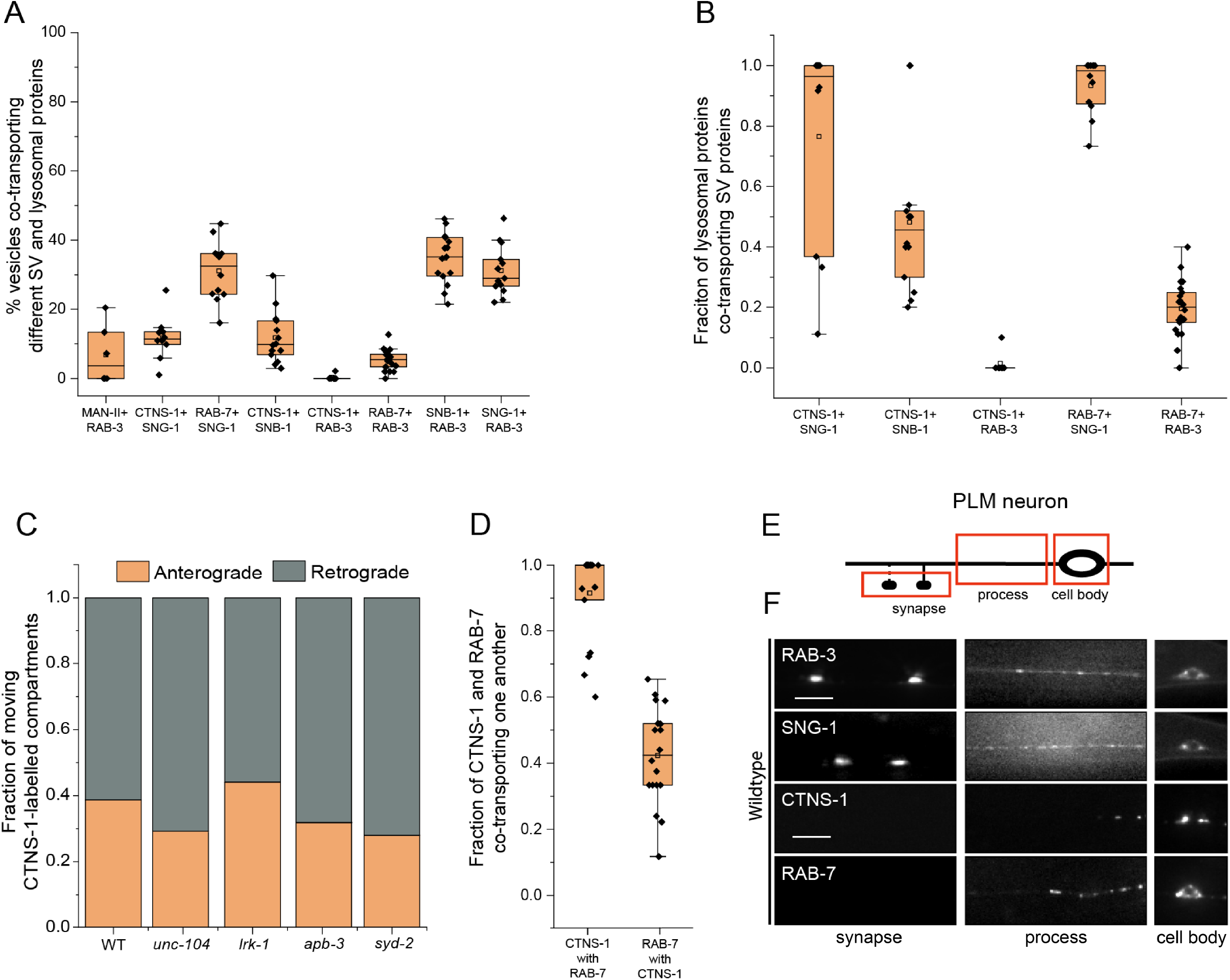
Synaptic vesicle proteins travel with lysosomal proteins in heterogenous carriers. (A) Quantitation of co-transport of different combinations of synaptic vesicle proteins and lysosomal proteins from kymograph analysis of dual color imaging. The number of animals per genotype (N) ≥ 10. Number of vesicles analyzed (n) > 600. (B) Quantitation of fraction of different lysosomal proteins co-transporting different synaptic vesicle proteins from kymograph analysis of dual color imaging. N ≥ 10; n > 100. (C) Quantitation of fraction of CTNS-1-labelled compartments moving in the anterograde and retrograde direction in different mutants. N ≥ 9 per genotype; the number of CTNS-1-labelled compartments ≥ 20. (D) Quantitation of co-transport of CTNS-1::mCherry and mNeonGreen::RAB-7 in WT animals from sequential dual color imaging at 1.3 fps. CTNS-1 with RAB-7 indicates the fraction of CTNS-1-labelled compartments co-transporting RAB-7. RAB-7 with CTNS-1 indicates the fraction of RAB-7-labelled compartments co-transporting CTNS-1. N ≥ 15 animals; n > 450. (E) Schematic showing the PLM neuron. Red boxes highlight the regions of imaging. The arrow shows the anterograde direction of vesicle motion. (F) GFP::RAB-3, SNG-1::GFP, CTNS-1::mCherry, and RAB-7::mScarlet in the cell body, process and synapses of wildtype PLM neurons. Scale bar: 10 μm.

To further characterize these compartments that contain both SVps and CTNS-1 or RAB-7, we examined their localization along the PLM process. Unlike SVps in wildtype animals, CTNS-1-labelled compartments are largely restricted to the PLM cell bodies, are present in the first 25 μm of the neuronal process in ∼27% of the animals, and never reach the PLM synapse (Fig. 1E, 1F, and S1H). RAB-7-labeled compartments are present in the proximal 25 μm of the neuronal process in 52% of the animals but are also absent from the PLM synapse (Fig. 1F and S1I).

Thus, lysosomal proteins that exit the cell body are present along with some SVps in carriers that we hereafter refer to as the SV-lysosomes or SV-lysosomal compartments. However, only a minority of SNG-1 or SNB-1 travel in CTNS-1-carrying compartments. RAB-3 is excluded from SV-lysosomal compartments and may therefore mark only SV precursors. For this study, we consider the CTNS-1-marked compartments as the SV-lysosomes, since nearly all CTNS-1-carrying compartments also contain SNG-1 (Fig. 1B).

### LRK-1 and the AP-3 complex exclude lysosomal proteins from SVp transport carriers

Mammalian AP-3 has been shown to play a role in separating lysosomal proteins from SVps (Newell-Litwa *et al*., 2009). As LRK-1 and APB-3 are also known to affect the trafficking of SVps (Choudhary *et al*., 2017), we investigated whether these genes regulate the trafficking of the SV-lysosome compartments in *C. elegans* TRNs.

There is a small but significant reduction in the co-transport of SNG-1 and RAB-3 in *lrk-1* (∼25%), but not in *apb-3* mutants (∼40%), which largely resembles the co-transport seen in wildtype TRNs (∼35%) (Fig. 2A, Suppl. Table 3, Movies S3 and S4). Interestingly, 45% of SNG-1-carrying vesicles co-transport CTNS-1 in *lrk-1*(*km17*), a kinase-deleted loss-of-function mutant. The frequency of SNG-1-carriers containing CTNS-1 is close to 60% in the *lrk-1* null, *lrk-1*(*km41*), and *apb-3* mutant animals (Fig. 2B and S2A, Suppl. Table 4). Nearly all CTNS-1-carrying compartments continue to transport SNG-1 (Fig. S2B; Suppl. Table 5). Furthermore, ∼80% and ∼65% of SNG-1-carrying vesicles co-transport RAB-7 in *lrk-1* and *apb-3* mutants, respectively (Fig. 2C and S2C; Suppl. Table 6, Movie S5). As with CTNS-1, nearly all RAB-7-carrying compartments continue to transport SNG-1 (Fig. S2D; Suppl. Table 7). Notably, the co-transport of CTNS-1 with SNB-1 is not affected in *lrk-1* and *apb-3* mutants (Fig. S2E; Suppl. Table 8), and RAB-3 continues to be absent from CTNS-1-carriers in these mutant animals (Fig. S2F; Suppl. Table 9). Thus, in both *lrk-1* and *apb-3* mutants, the SV-lysosomal compartments show significant transport into the neuronal process, suggesting that both LRK-1 and the AP-3 complex play key roles in sorting CTNS-1 and RAB-7 away from SVps. In these mutants, RAB-3 continues to be excluded from SV-lysosomal compartments and RAB-3 may, therefore, mark the only SVp-containing carriers.

**Figure 2:**
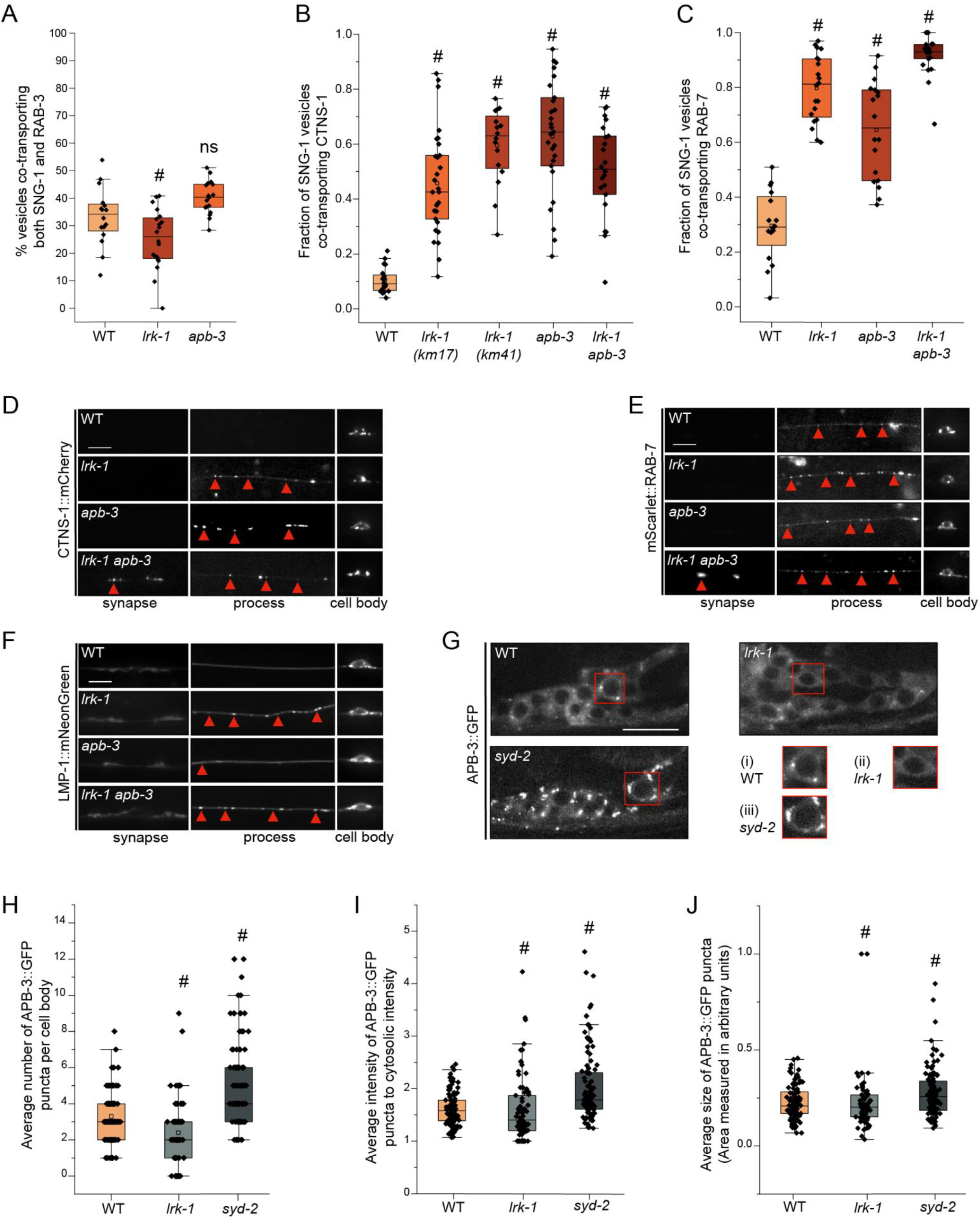
LRK-1 and AP-3 act in parallel and through SYD-2 to regulate lysosomal protein trafficking. (A) Quantitation of co-transport of SNG-1::eGFP and mCherry::RAB-3 in WT, *lrk-1*(*km17*), and *apb-3*(*ok429*) from kymograph analysis of sequential dual color imaging at 1.3 frames per second (fps). # P-value ≤ 0.05 (One-Way ANOVA with Tukey’s post-hoc test, all comparisons to WT); ns: not significant; Number of animals (N) ≥ 15 per genotype; Number of vesicles analyzed per genotype (n) > 800. (B) Quantitation of fraction of SNG-1-carrying vesicles co-transporting CTNS-1 in WT, *lrk-1*(*km17*), *lrk-1*(*km41*), *apb-3*(*ok429*) and *lrk-1*(*km17*) *apb-3*(*ok429*), from kymograph analysis of sequential dual color imaging at 1.3 fps. # P-value ≤ 0.05 (Mann–Whitney Test); N ≥ 15 per genotype; n > 500. (C) Quantitation of fraction of SNG-1-carrying vesicles co-transporting RAB-7 from WT, *lrk-1*(*km17*), *apb-3*(*ok429*) and *lrk-1*(*km17*) *apb-3*(*ok429*), kymograph analysis of dual color imaging. # P-value ≤ 0.05 (One-Way ANOVA with Tukey’s post-hoc test, all comparisons to WT); ns: not significant; N ≥ 20 per genotype; n > 800. (D) CTNS-1::mCherry in the cell body, process, and synapses of PLM neurons of WT, *lrk-1*(*km17*), *apb-3*(*ok429*), and *lrk-1*(*km17*) *apb-3*(*ok429*). Scale bar: 10 μm. Red arrows point to some CTNS-1-labelled compartments. (E) mScarlet::RAB-7 in the cell body, process, and synapses of PLM neurons of WT, *lrk-1*(*km17*), *apb-3*(*ok429*), and *lrk-1*(*km17*) *apb-3*(*ok429*). Scale bar: 10 μm. Red arrows point to some RAB-7-labelled compartments. (F) LMP-1::mNeonGreen in the cell body, process, and synapses of PLM neurons of WT, *lrk-1*(*km17*), *apb-3*(*ok429*), and *lrk-1*(*km17*) *apb-3*(*ok429*). Scale bar: 10 μm. Red arrows point to some RAB-7-labelled compartments. (G) Images showing APB-3::GFP puncta in the head ganglion cell bodies of WT, *lrk-1*(*km17*), and *syd-2*(*ok217*). Scale bar: 10 μm. Red boxes highlight the regions of insets with cell bodies from images showing APB-3::GFP in (i) WT, (ii) *lrk-1*, and (iii) *syd-2*. (H) Quantitation of the number of APB-3::GFP puncta per cell body in WT, *lrk-1*(*km17*), and *syd-2*(*ok217*). # P-value ≤ 0.05 (Mann–Whitney Test); ns: not significant; N > 10 animals; n > 75 cell bodies. (I) Quantitation of intensity of APB-3::GFP puncta in cell bodies of WT, *lrk-1*(*km17*), and *syd-2*(*ok217*). The ratio of the intensity of APB-3::GFP puncta to cytosolic intensity in the cell body is plotted. # P-value ≤ 0.05 (Mann–Whitney Test); ns: not significant; N > 10 animals; n > 75 cell bodies. (J) Quantitation of average size of APB-3::GFP puncta per cell body in WT, *lrk-1*(*km17*), and *syd-2*(*ok217*). # P-value ≤ 0.05 (Mann–Whitney Test); ns: not significant; N > 10 animals; n > 75 cell bodies.

The localization of lysosomal proteins CTNS-1 and RAB-7 is also altered in both *lrk-1* and *apb-3* mutants. In contrast to wildtype, CTNS-1 is localized to the first 25 μm of the PLM neuronal process in ∼63% of *lrk-1* and ∼45% of *apb-3* mutants (Fig. 2D, S1H), while 100% of *lrk-1* and ∼80% of *apb-3* mutants show RAB-7 up to 25 μm away from the cell body (Fig. 2E and S1I). Like CTNS-1 and RAB-7, LMP-1 is largely restricted to the cell body in wildtype animals (Fig. 2F). However, unlike CTNS-1 and RAB-7, LMP-1 localization is only affected in *lrk-1* mutants, with LMP-1 localizing along the neuronal process until the first 50 μm in all *lrk-1* animals. In contrast to *lrk-1*, LMP-1 does not localize beyond the first 25 μm of the PLM neuron in *apb-3* mutant animals (Fig. 2F, S1J, and S1K). Since both LMP-1 localization and co-transport of SNG-1 and RAB-3 are affected only in *lrk-1* mutants, LRK-1 likely affects the trafficking of more kinds of SVp carriers than the AP-3 complex.

The *lrk-1 apb-3* double mutants, similar to *lrk-1* single mutants, show a significant increase in the co-transport of CTNS-1 and RAB-7 with SNG-1 (Fig. 2B and 2C, Suppl. Tables 4 and 6). However, there is an increased number of *lrk-1 apb-3* double mutant animals with CTNS-1 localized along the neuronal process than in either *lrk-1* or *apb-3* single mutants (Fig. 2D and S1H). The *lrk-1 apb-3* double mutants show a similar frequency of animals with RAB-7 and LMP-1 localized along the neuronal process as seen in *lrk-1* mutant (Fig. 2E, 2F, S1I, S1J, and S1K). These data suggest that LRK-1 may act upstream of APB-3 in the trafficking of SV-lysosome compartments.

### LRK-1 regulates localization of the AP-3 complex

LRK-1 acts via the AP-1 and the AP-3 complexes to regulate polarized SVp trafficking and the trafficking of SVp transport carriers. LRK-1 is known to assist in the Golgi membrane localization of the AP-1 clathrin adaptor complex, thereby regulating its function (Choudhary *et al*., 2017). To examine whether LRK-1 regulates the membrane localization of the AP-3 complex as well, we examined the distribution of the β subunit of the AP-3 complex, APB-3::GFP, in neuronal cell bodies of *lrk-1* mutants (Fig. 2G). In wildtype, APB-3::GFP shows punctate localization in the cell body with an average of ∼2 to 4 puncta/cell body. In *lrk-1* mutant animals, there are fewer APB-3::GFP puncta per cell body and more cell bodies that lack puncta (Fig. 2G (i) and (ii), 2H and S2G; Suppl. Table 10). The average intensity and size of the APB-3::GFP puncta remain largely unaltered in *lrk-1* mutants (Fig. 2I and 2J; Suppl. Tables 11 and 12). This suggests that, as observed with the AP-1 complex (Choudhary *et al*., 2017), in *lrk-1* mutants the AP-3 complex may not be recruited efficiently to membrane surfaces. Some of the sorting roles of LRK-1 are likely mediated by facilitating AP-3 localization to membrane surfaces.

### SV-lysosomes in *lrk-1* and *apb-3* mutants are dependent on UNC-104

SVps are known to be dependent on the anterograde motor UNC-104/KIF1A for their exit from neuronal cell bodies (Hall and Hedgecock, 1991; Pack-Chung *et al*., 2007; Okada and Hirokawa, 1999; Kumar *et al*., 2010). SV-lysosomes likely depend on IMAC/KIF1A in *Drosophila* neurons (Vukoja *et al*., 2018). However, the transport of SVp carriers in *lrk-1* and *apb-3* mutants is only partially dependent on UNC-104 (Choudhary *et al*., 2017). To examine the UNC-104 dependence of the SV-lysosomes, we characterized the role of UNC-104 in transporting both SVs and SV-lysosomes out of neuronal cell bodies using a weak cargo binding-defective hypomorph, *unc-104*(*e1265tb120*) (Kumar *et al*., 2010).

SNG-1 in *lrk-1* is partially dependent on UNC-104, as little SNG-1 reaches the synapse in *lrk-1*; *unc-104* compared to that in *lrk-1* mutant animals (Fig. 3B). In a strong loss-of-function *unc-104* allele, RAB-3 in *lrk-1* is partially dependent on UNC-104 (Choudhary *et al*., 2017). However, with a weak loss-of-function *unc-104* allele, RAB-3 reaches the synapse in *lrk-1*; *unc-104* double mutants (Fig. S3A). SNG-1- and RAB-3-carriers in *apb-3* are partially dependent on UNC-104, as both markers do not reach the synapse in *apb-3; unc-104* double mutants (Fig. 3B and S3A). This suggests that SVps in *lrk-1* and *apb-3* mutants are only partially dependent on UNC-104. Additionally, RAB-3 and SNG-1 in these mutants appear to have slightly different extents of dependence on UNC-104.

**Figure 3:**
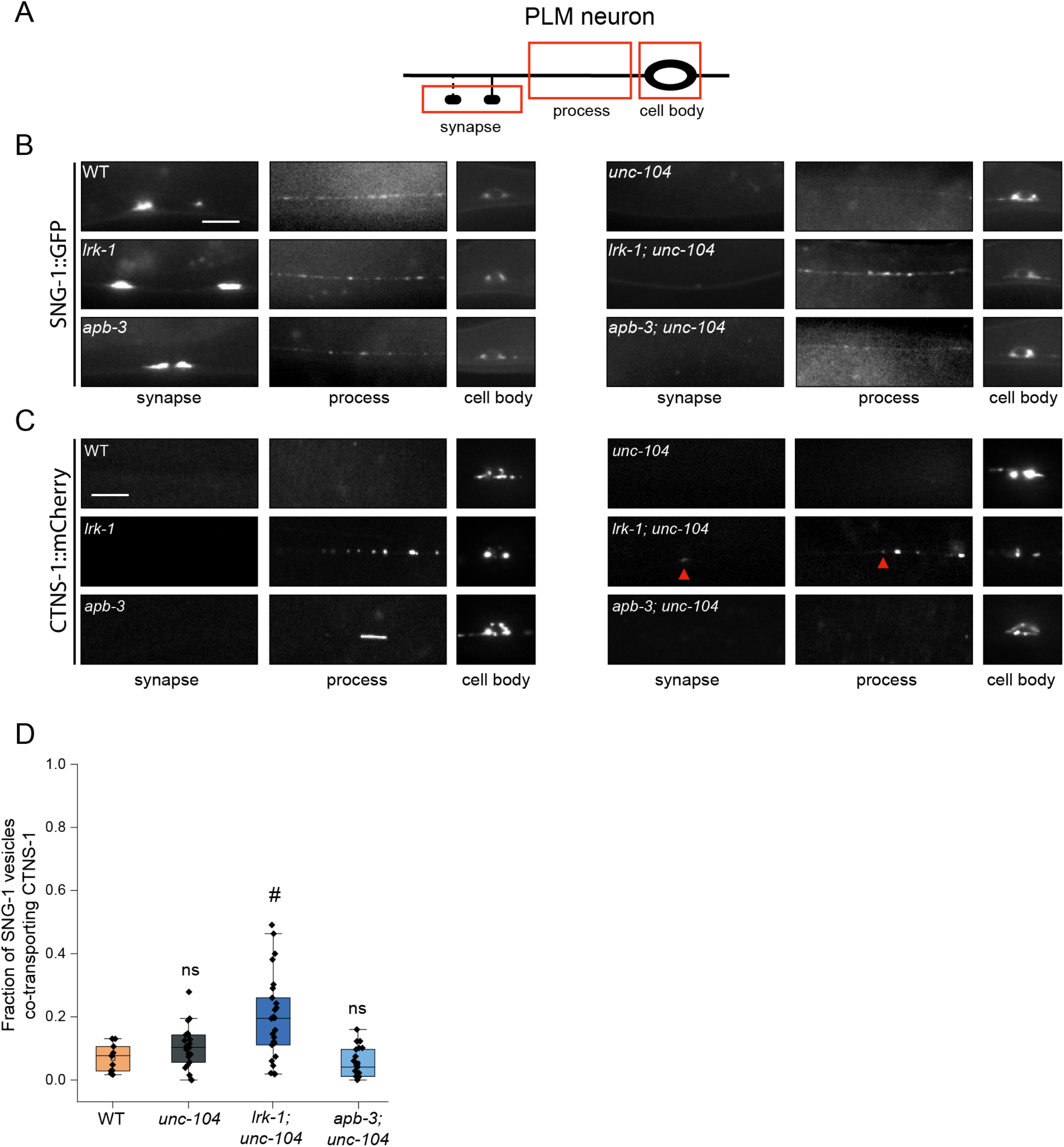
SV-lysosomes in *lrk-1* and *apb-3* mutants are dependent on UNC-104. (A) Schematic showing the PLM neuron. Red boxes highlight the regions of imaging. (B) SNG-1::GFP in the cell body, process, and synapses of PLM neurons showing dependence on UNC-104 in *lrk-1*(*km17*) and *apb-3*(*ok429*) mutants and their doubles with *unc-104*(*e1265tb120*). Scale bar: 10 μm. (C) CTNS-1::mCherry in the cell body, process, and synapses of PLM neurons showing dependence on UNC-104 in *lrk-1*(*km17*) and *apb-3*(*ok429*) mutants and their doubles with *unc-104*(*e1265tb120*). Red arrows highlight fainter CTNS-1 puncta. Scale bar: 10 μm. (D) Quantitation of co-transport of SNG-1 and CTNS-1 in *unc-104*(*e1265tb120*), *lrk-1*(*km17*); *unc-104*, and *apb-3*(*ok429*); *unc-104* from kymograph analysis of sequential dual color imaging done at 1.3 fps. #P-value ≤ 0.05 (Mann–Whitney Test, all comparisons to WT); ns: not significant; Number of animals (N) ≥ 18 per genotype; Number of vesicles (n) > 1200.

We next examined the CTNS-1-containing SV-lysosomes, which are dependent on UNC-104 for their exit from the cell body (Fig. 3C, S3B, S1H). The localization of CTNS-1 is dependent on UNC-104 in *apb-3* mutants but not in *lrk-1* mutants (Fig. 3C and S1H). Compared to *lrk-1* single mutants, there is an increased number of animals with CTNS-1 localized along the neuronal process in *lrk-1; unc-104* double mutants (Fig. 3C and S1H). The fraction of SNG-1-carrying vesicles co-transporting CTNS-1 is comparable in wildtype, *unc-104*, and *apb-3; unc-104* animals (Fig. 3D; Suppl. Table 13, movie S6). However, in *lrk-1; unc-104* mutants, a higher number of SNG-1-carrying vesicles co-transport CTNS-1 (21%), but this is lower than that observed in *lrk-1* mutants alone (45%) (Fig. 2B, 3D).

These data suggest that the axonal localization of SV-lysosomal compartments is dependent on UNC-104 in wildtype and *apb-3* mutants but independent of UNC-104 in *lrk-1* mutants. The extent of localization along the neuronal process in *apb-3* mutants is likely to reflect the net transport activity of UNC-104. The SV-lysosomes in *lrk-1* likely depend on other motors for their transport into axons, and in the absence of UNC-104, these alternate motors may allow more SV-lysosomes to localize along the neuronal process. Additionally, UNC-104 may play an indirect role in the sorting of CTNS-1 from SNG-1 compartments in *lrk-1* mutants.

### SV-lysosomes in *lrk-1* and *apb-3* mutants are differentially dependent on SYD-2

The active zone protein SYD-2 has been shown to regulate lysosomal protein distribution in *C*. *elegans* neurons (Edwards *et al*., 2015a). SYD-2 is also a known genetic enhancer of UNC-104 and is known to directly bind this motor (Zheng *et al*., 2014; Shin *et al*., 2003; Wagner *et al*., 2009; Stucchi *et al*., 2018). We, therefore, examined whether the altered localization of the SV-lysosomal compartments in *lrk-1* and *apb-3* depends on SYD-2. We used two different alleles of *syd-2*, the null allele, *syd-2*(*ok217*), and the loss-of-function allele, *syd-2*(*ju37*), with a premature stop codon in the LH2 domain (Zhen and Jin, 1999; Wagner *et al*., 2009). The N-terminal portion of SYD-2 expressed in *syd-2*(*ju37*) allele is capable of physically associating with UNC-104 (Wagner *et al*., 2009).

Both *syd-2(ok217)* and *syd-2*(*ju37*) resemble wildtype in the number of SNG-1-carriers co-transporting CTNS-1 as well as in the number of animals showing CTNS-1 localization along the PLM neuronal process (Fig. 4A and S1H, Movie S7). The co-transport of CTNS-1 in SNG-1-carrying carriers is similar (∼55–60%) in *lrk-1*, *apb-3*, and *lrk-1; syd-2(ok217)* mutant animals (Fig. 4A; Suppl. Table 14). Similarly, the number of SNG-1-carriers co-transporting RAB-7 in TRNs is comparable (∼90%) in *lrk-1* and *lrk-1; syd-2(ok217)* mutant animals (Fig. 4B; Suppl. Table 15). Additionally, both *lrk-1; syd-2*(*ok217*) and *lrk-1; syd-2*(*ju37*) show an increased or similar number of animals in which CTNS-1 or RAB-7 are localized along the neuronal process compared to *lrk-1* alone (Fig. 4C, 4D, 4E and S1H). The number of animals showing LMP-1 in the neuronal process in *lrk-1* and *lrk-1; syd-2*(*ok217*) is similar (Fig. 4F, S1J, and S1K). Thus, SYD-2 does not appear to be required for the transport or localization of SV-lysosomes. In the absence of SYD-2, more *lrk-1* animals show a greater number of SV-lysosomes along the neuronal process, akin to the phenotypes observed in *lrk-1; unc-104*, suggesting that SYD-2 might function like UNC-104.

**Figure 4:**
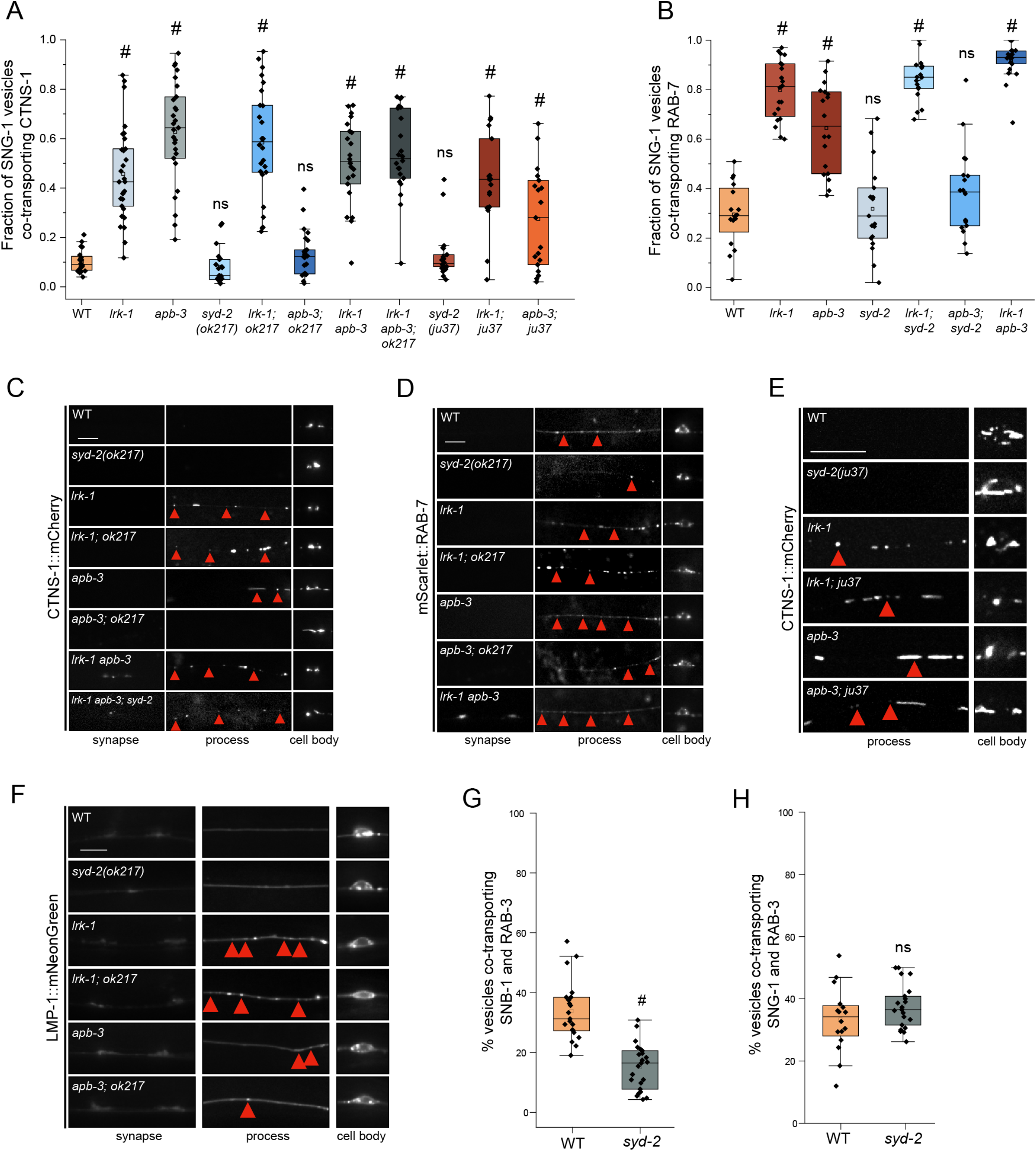
Distribution of SV-lysosomal compartments depends on UNC-104. (A) Quantitation of co-transport of SNG-1 and CTNS-1 in *syd-2* mutants and their doubles with *lrk-1*(*km17*) and *apb-3*(*ok429*), from kymograph analysis of dual color imaging. *ok217* refers to the null allele of *syd-2*, *syd-2*(*ok217*); while *ju37* refers to the *syd-2*(*ju37*) allele. #P-value ≤ 0.05 (Mann–Whitney Test, all comparisons to WT); ns: not significant; Number of animals (N) ≥ 18 per genotype; Number of vesicles (n) > 750. Values for *lrk-1* and *apb-3* single mutants are the same as those in Fig. 2B. (B) Quantitation of co-transport of SNG-1 and RAB-7 in *syd-2*(*ok217*) and its doubles with *lrk-1*(*km17*) and *apb-3*(*ok429*), from kymograph analysis of dual color sequential imaging at 1.3 fps. P-value > 0.05 (One-Way ANOVA with Tukey’s post-hoc test); ns: not significant; N ≥ 19 per genotype; n > 700. Values for *lrk-1* and *apb-3* single mutants are the same as those in Fig. 2C. (C) CTNS-1::mCherry in the cell body, process, and synapses of PLM neurons of *syd-2*(*ok217*) mutant and its doubles with *lrk-1*(*km17*) and *apb-3*(*ok429*). Red arrows highlight some CTNS-1-carrying compartments, some fainter. Scale bar: 10 μm. (D) mScarlet::RAB-7 in the cell body, process, and synapses of PLM neurons of *syd-2*(*ok217*) mutant and its doubles with *lrk-1*(*km17*) and *apb-3*(*ok429*). Red arrows highlight some RAB-7-carrying compartments, some fainter. Scale bar: 10 μm. (E) CTNS-1::mCherry in the cell body, process, and synapses of PLM neurons of *syd-2*(*ju37*) mutant and its doubles with *lrk-1*(*km17*) and *apb-3*(*ok429*). Red arrows highlight some CTNS-1-carrying compartments, some fainter. Scale bar: 10 μm. Imaged at 100×. (F) LMP-1::mNeonGreen in the cell body, process, and synapse of PLM neurons of *syd-2*(*ok217*) mutant and its doubles with *lrk-1*(*km17*) and *apb-3*(*ok429*). Red arrows indicate LMP-1-carrying compartments. Scale bar: 10 μm. (G) Quantitation of co-transport of SNB-1 and RAB-3, in *syd-2*(*ok217*), from simultaneous dual color imaging at 3 frames per second (fps). # P-value ≤ 0.05 (One-Way ANOVA with Tukey’s post-hoc test); N > 20. (H) Quantitation of co-transport of SNG-1 and RAB-3, in *syd-2*(*ok217*), from sequential dual color imaging at 1.3 fps. P-value > 0.05 (One-Way ANOVA with Tukey’s post-hoc test); ns: not significant; N > 15.

In contrast to the above phenotypes observed with *lrk-1*, the number of SNG-1 carriers co-transporting CTNS-1 or RAB-7 in *apb-3; syd-2(ok217)* (∼40%) is similar to that in wildtype, and is lower than that seen in *apb-3* mutants alone (Fig. 4A and 4B; Suppl. Tables 11 and 12). The number of animals with CTNS-1 and RAB-7 localized along the neuronal process is lower in *apb-3; syd-2(ok217)* than in *apb-3* mutants (Fig. 4C, 4D, and S1H). Unlike these markers, the number of LMP-1-marked carriers in the neuronal process of *apb-3; syd-2(ok217)* is increased compared to that in *apb-3* mutants (Fig. 4F, S1J, and S1K). Further, unlike in *apb-3; syd-2*(*ok217*), the *apb-3; syd-2*(*ju37*) animals show increased co-transport of CTNS-1 with SNG-1 (30%) compared to wildtype (10%) but lower than that in *apb-3* mutants (63%) (Fig. 4A, Suppl. Tables 14 and 15). Furthermore, *lrk-1; syd-2*(*ju37*) and *apb-3; syd-2*(*ju37*) show an increased number of animals with CTNS-1 localized along the neuronal process compared to *lrk-1* and *apb-3* mutants, respectively (Fig. 4E and S1H). As with *lrk-1* and *apb-3* mutants, the co-transport of CTNS-1 with either SNB-1 or RAB-3 is unaffected in *syd-2(ok217)* (Fig. S3C and S3D; Suppl. Tables 16 and 17). As the *apb-3* phenotypes appear to be dependent on the presence of SYD-2, it is likely that *syd-2* acts downstream of *apb-3*. The genetic interaction of *apb-3* with the two *syd-2* alleles suggests that the N-terminal region of SYD-2, which is known to bind UNC-104/KIF1A, is sufficient to enable the exit of SV-lysosomes in the neuronal processes of *apb-3* mutants.

We next examined whether *syd-2* affects the composition of the SVp carrier pools without lysosomal proteins. The incidence of co-transport of SNB-1 and RAB-3 is lower in *syd-2*(*ok217*) mutants (∼15%) than in wildtype (∼35%) (Fig. 4G; Suppl. Table 18), similar to that reported in *lrk-1* and *apb-3* single mutants (Choudhary *et al*., 2017). Furthermore, *syd-2(ok217)*, like *apb-3*, does not affect the co-transport of SNG-1 and RAB-3 (Fig. 4H; Suppl. Table 19). These observations suggest that SYD-2 only affects the trafficking of a subset of SVps and may act downstream of APB-3.

In *syd-2(ok217)*, the number of APB-3::GFP puncta per cell body increase (Fig. 2G, 2H and S2G), while the size and average intensity of these puncta remain comparable to wildtype animals (Fig. 2G(iii), 2I and 2J; Suppl. Tables 11 and 12). This suggests that *syd-2* may not influence the ability of the AP-3 complex to associate with membrane surfaces, but rather acts on the compartments formed after AP-3 has acted on them.

To further determine the hierarchy of action of SYD-2 on compartments formed in *lrk-1* and *apb-3*, we examined *lrk-1 apb-3*; *syd-2* triple mutants. Notably, *lrk-1 apb-3*; *syd-2* triple mutants are similar to *lrk-1 apb-3* in the co-transport of CTNS-1 with SNG-1 (Fig. 4A, Suppl. Table 14). Furthermore, both *lrk-1 apb-3* double mutants and *lrk-1 apb-3*; *syd-2* triple mutants show a similar number of animals with CTNS-1 compartments localized along the neuronal process (Fig. 4A, 4C, and S1H). These data are consistent with a hierarchical pathway wherein LRK-1 acts upstream of the AP-3 complex, and SYD-2 acts downstream of AP-3 to facilitate UNC-104 activity.

### UNC-104 and SYD-2 are necessary for SV-lysosome transport in *apb-3* mutants

Previous studies have shown that the N-terminal region of Liprin-α/SYD-2 binds to KIF1A/UNC-104 (Shin *et al*., 2003; Wagner *et al*., 2009; Stucchi *et al*., 2018). This physical interaction between SYD-2 and UNC-104, in addition to the genetic interactions that we and others observe, suggests that SYD-2 may act through UNC-104 to facilitate both motor and its cargo’s transport (Wagner *et al*., 2009; Zheng *et al*., 2014). Further, we have shown that SVp carriers in mutants of *lrk-1* and *apb-3* are only partially dependent on UNC-104 for their transport (Choudhary *et al*., 2017). Therefore, we examined the potential role of the UNC-104–SYD-2 complex in the localization and transport of SVp carriers and SV-lysosomes.

In wildtype animals, the localization of the transmembrane SVp SNG-1 is dependent on UNC-104 but not on SYD-2 (Fig. 3B and 5B). However, *unc-104*; *syd-2* mutants have less SNG-1 in the PLM neuronal process compared to that seen in *unc-104* single mutants, demonstrating that SYD-2 facilitates UNC-104-dependent SVp transport (Fig. 5B) (Zheng *et al*., 2014). Transport of SVps is not dependent on SYD-2 in either *lrk-1* or *apb-3* mutants (Fig. 5B). *lrk-1*; *unc-104* and *lrk-1*; *unc-104*; *syd-2* triple mutants show comparable SNG-1 localization in the neuronal process. Likewise, *apb-3*; *unc-104* and *apb-3*; *unc-104*; *syd-2* also show comparable SNG-1 localization along the neuronal process (Fig. 5B). However, in *lrk-1*; *unc-104*; *syd-2* triple mutants, the peripherally-associated membrane protein RAB-3 does not reach the synapse unlike in *lrk-1*; *unc-104* double mutants (Fig. S3A). Unlike the phenotypes with *lrk-1*, RAB-3 localization in *apb-3*; *unc-104*; *syd-2* and *apb-3*; *unc-104* is similar (Fig. S3A). Together, these data indicate that SYD-2 does not facilitate UNC-104-dependent SVp transport in *apb-3* mutants.

**Figure 5:**
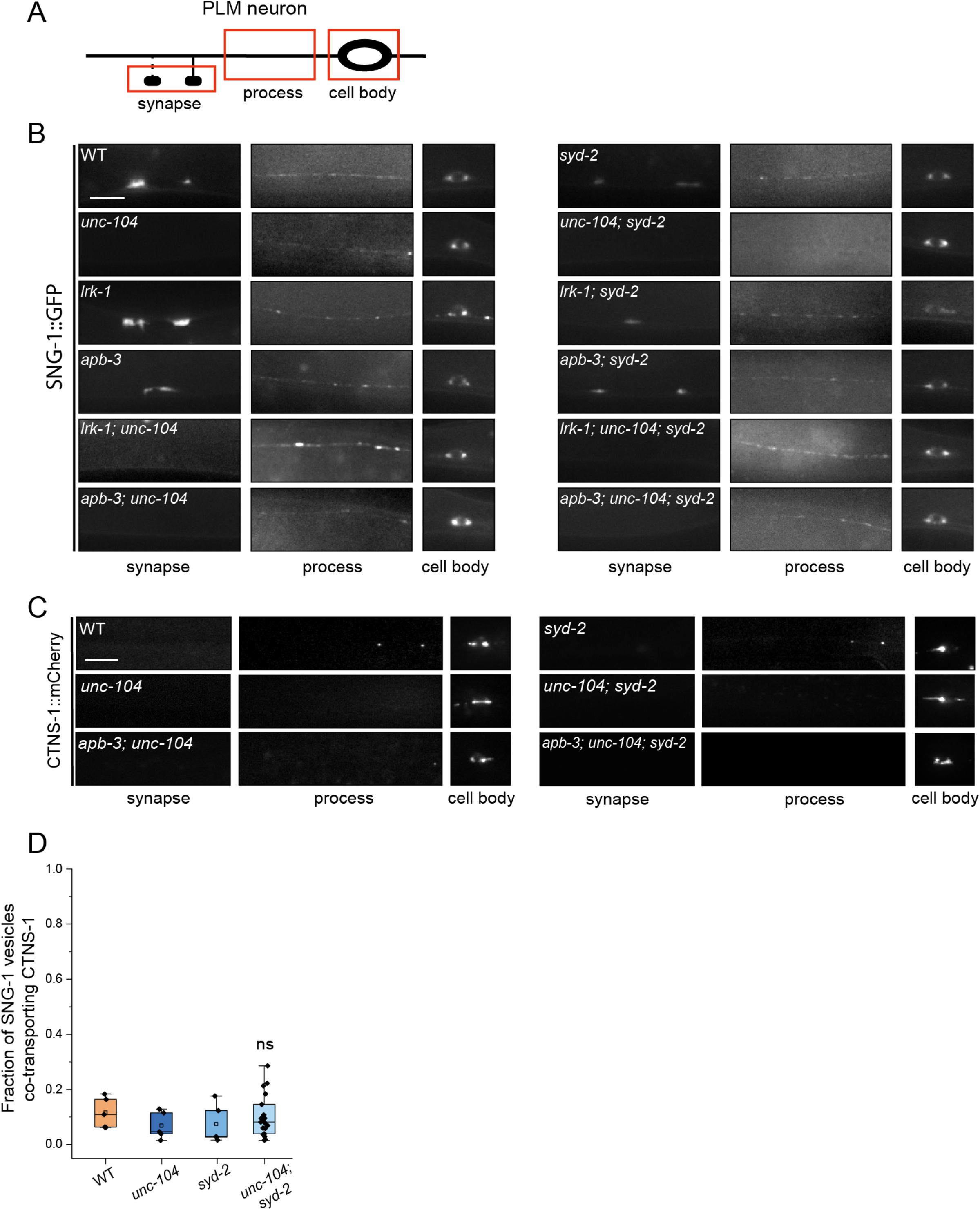
SYD-2 is required for UNC-104 dependent of SVp carriers. (A) Schematic of the PLM neuron. The red box highlights the region of imaging. (B) SNG-1::GFP in the cell body, process and synapses of PLM neurons of *syd-2*(*ok217*) and *unc-104*(*e1265tb120*), and their doubles with *lrk-1*(*km17*) and *apb-3*(*ok429*). Scale bar: 10 μm. (C) CTNS-1::mCherry in the cell body, process and synapses of PLM neurons of *syd-2*(*ok217*) and *unc-104*(*e1265tb120*), and their doubles with *apb-3*(*ok429*). Scale bar: 10 μm. (D) Quantitation of co-transport of SNG-1 and CTNS-1 in WT, *unc-104*(*e1265tb120*), *syd-2*(*ok217*), and *unc-104*; *syd-2* from kymograph analysis of sequential dual color imaging at 1.3 fps. P-value > 0.05 (Mann–Whitney Test, all comparisons to WT); ns: not significant; Number of animals (N) > 20 for *unc-104*; *syd-2*; Number of vesicles (n) >1000.

SV-lysosomes, marked by CTNS-1, are dependent on UNC-104 and largely independent of SYD-2 in wildtype (Fig. S3B). The number of SNG-1 vesicles co-transporting CTNS-1 is similar in *unc-104*, *syd-2*, and *unc-104*; *syd-2* mutant animals (Fig. 5D; Suppl. Table 20), suggesting that these genes do not regulate the sorting of lysosomal and SV proteins away from each other.

In *lrk-1* mutants, SV-lysosomes are independent of SYD-2 and UNC-104 (Fig. 3C). *lrk-1*; *unc-104* and *lrk-1*; *unc-104*; *syd-2* triple mutants show comparable localization of CTNS-1 along the neuronal process (Fig. S1H), suggesting that SYD-2 does not facilitate UNC-104-dependent SV-lysosome trafficking in *lrk-1* mutants. The CTNS-1-marked SV-lysosomes in *apb-3* mutants are dependent on both UNC-104 and SYD-2 (Fig. 3C and 5C). The *apb-3*; *unc-104*; *syd-2* triple mutants are similar to the *apb-3; unc-104* and *apb-3; syd-2* mutants (Fig. 5C and S1H). These data suggest that SYD-2 is important for the localization of the SV-lysosomal compartments along the neuronal process in *apb-3* mutants.

SVp carriers depend on both UNC-104 and SYD-2 in wildtype, but the SNG-1-containing compartment only partially depends on UNC-104 but not SYD-2 in both *lrk-1* and *apb-3* mutants. SV-lysosomes depend on UNC-104 but appear to be largely independent of SYD-2 in wildtype. In *lrk-1* mutants, the SV-lysosomes appear to be independent of both UNC-104 and SYD-2. In *apb-3* mutants, SV-lysosomes are dependent on both UNC-104 and SYD-2. This suggests that, in *apb-3* mutants, the preference for the UNC-104-SYD-2 complex is switched between the SVp alone-containing compartments (both SNG-1 and RAB-3) and the SV-lysosomes. The action of AP-3 appears essential to ensure that SYD-2 facilitates UNC-104-dependent transport of SVps.

### SYD-2 and the AP-1 complex together regulate the polarized distribution of SVps

SNB-1-labeled SVp carriers in *lrk-1* and *apb-3* have been shown to mislocalize to the dendrites (Sakaguchi-Nakashima *et al*., 2007; Choudhary *et al*., 2017). Since *syd-2* phenocopies *lrk-1* and *apb-3* in affecting the co-transport of SNB-1 and RAB-3 (Fig. 4G) (Choudhary *et al*., 2017), we examined whether SVps mislocalize to dendrites in *syd-2* mutants. Similar to wildtype, SNB-1 was found to be excluded from the dendrites of the ASI neuron in *syd-2*(*ok217*) (Fig. 6B, Table 1), which shows a similar orientation of axonal and dendritic microtubules as wildtype (Fig. S4A). Thus, SYD-2 does not appear to play a key role by itself in regulating polarized trafficking of SVps.

**Figure 6:**
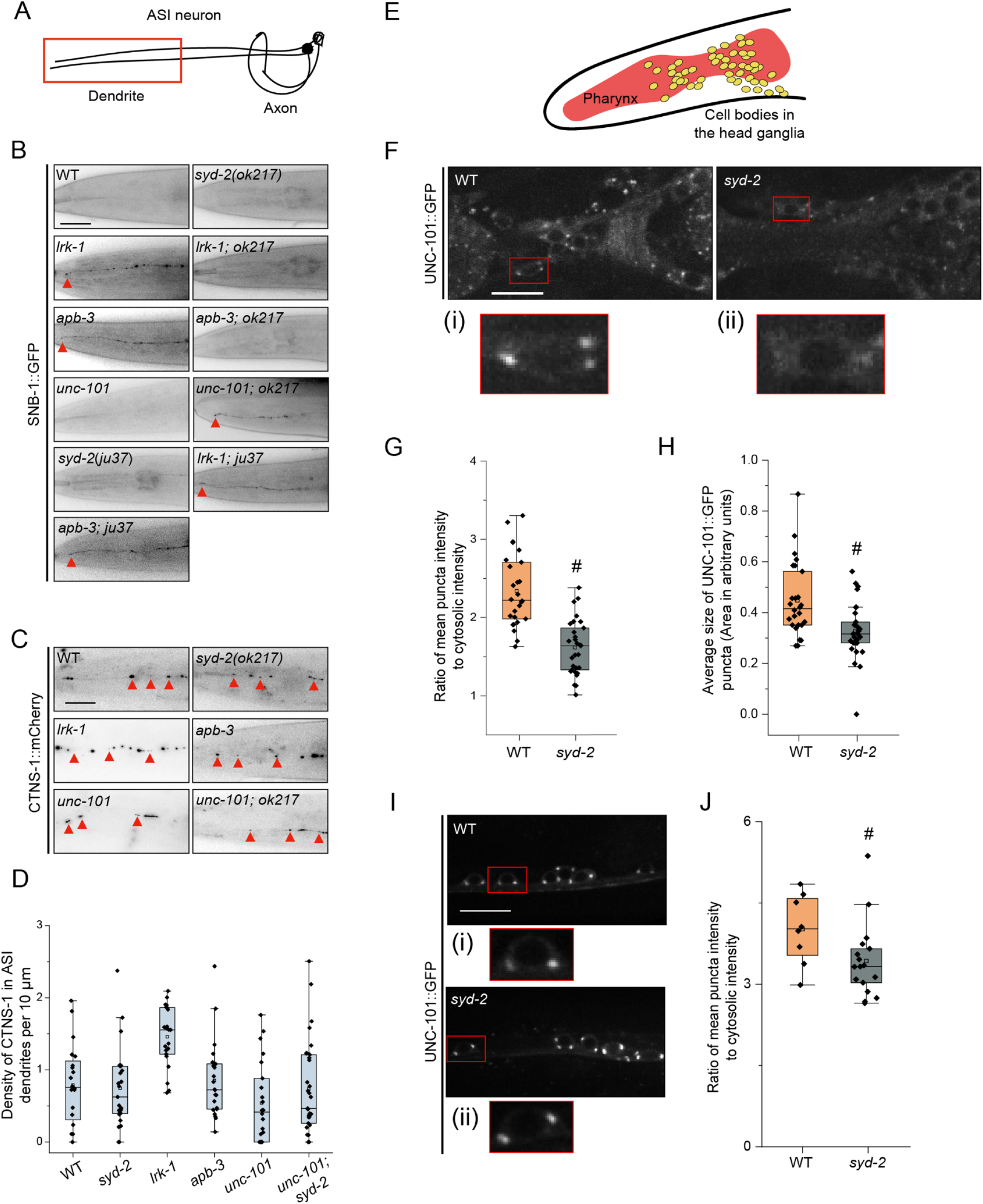
SYD-2 and the AP-1 complex together regulate the polarized distribution of SVps to axons. (A) Schematic of the ASI chemosensory neuron. Red box highlights the region of imaging. (B) SNB-1::GFP in the dendrite of the ASI neuron of WT and two alleles of *syd-2* and their doubles with *lrk-1*(*km17*) and *apb-3*(*ok429*). *ok217* represents *syd-2*(*ok217*) allele. *ju37* represents *syd-2*(*ju37*) allele. *unc-101*(*m1*) is a substitution mutation in the μ chain of the AP-1 complex causing a premature stop. Red arrows point to the SNB-1::GFP signal at the dendrite tip. Scale bar: 20 μm. Number of animals (N) > 6 per all single mutant genotypes; N > 20 for all double mutant genotypes. (C) CTNS-1::mCherry in the dendrite of the ASI neurons of WT, *syd-2*(*ok217*), *lrk-1*(*km17*), *apb-3*(*ok429*), *unc-101*(*m1*), and *unc-101*(*m1*); *syd-2*(*ok217*). Red arrows point to CTNS-1 compartments in the dendrite. Scale bar: 20 μm. N > 20 per genotype. (D) Density (number of CTNS-1 puncta per 10 μm in the ASI dendrite) of CTNS-1 in the ASI dendrite. # P-values ≤ 0.05 (Mann–Whitney Test, black comparisons against WT and blue comparisons against *lrk-1*); N > 20 for each genotype. (E) Schematic of *C*. *elegans* head showing the pharynx (red) and the head ganglion cell bodies (yellow). (F) Images showing UNC-101::GFP puncta in the head ganglion cell bodies of WT and *syd-2*(*ok217*). Scale bar: 10 μm. The red boxes highlight the regions of insets with cell bodies from images showing UNC-101::GFP in (i) WT and (ii) *syd-2*. (G) Quantitation of intensity of UNC-101::GFP puncta in the head ganglion cell bodies in WT and *syd-2*(*ok217*). The ratio of the intensity of UNC-101::GFP puncta to cytosolic intensity in the cell body is plotted. # P-value ≤ 0.05 (One-Way ANOVA with Tukey’s post-hoc test); N > 5 animals; n > 25 cell bodies. (H) Quantitation of average size of UNC-101::GFP puncta per cell body in WT and *syd-2*(*ok217*). # P-value ≤ 0.05 (Mann–Whitney Test); N > 5 animals; n > 25 cell bodies. (I) Images showing UNC-101::GFP puncta in the cell bodies of the ventral nerve cord neurons in WT and *syd-2*(*ok217*). Scale bar: 10 μm. The red boxes highlight the regions of insets with cell bodies from images showing UNC-101::GFP in (i) WT and (ii) *syd-2*. (J) Quantitation of intensity of UNC-101::GFP puncta in the cell bodies of the ventral nerve cord in WT and *syd-2*(*ok217*). The ratio of the intensity of UNC-101::GFP puncta to cytosolic intensity in the cell body is plotted. # P-value ≤ 0.05 (Mann–Whitney test); N > 5 animals; n > 10 cell bodies.

**Table 1:** SNB-1 in ASI

| **Genotype** | **Average % length of dendrite showing SNB-1 signal** | **Standard deviation** |
| --- | --- | --- |
| WT | 39 % | 27 |
| *lrk-1(km17)* | 92 % | 2 |
| *apb-3(ok429)* | 69 % | 28 |
| *syd-2(ok217)* | 25 % | 25 |
| *lrk-1(km17); syd-2(ok217)* | 35 % | 23 |
| *apb-3(ok429); syd-2(ok217)* | 45 % | 28 |

Previous studies have shown that the dendritic mislocalization of SNB-1 in *lrk-1* depends on the UNC-101/µ subunit of the AP-1 complex (Sakaguchi-Nakashima *et al*., 2007; Choudhary *et al*., 2017). Therefore, we examined whether *syd-2* genetically interacts with *unc-101* to regulate the polarized distribution of SVps. SNB-1 is absent from dendrites in *syd-2* and *unc-101* single mutant animals, while *unc-101*; *syd-2*(*ok217*) fails to exclude SNB-1 from the ASI dendrite (Fig. 6B). Further, SNB-1 dendritic mislocalization is suppressed in *lrk-1*; *syd-2*(*ok217*) and *apb-3*; *syd-2*(*ok217*) mutants (Fig. 6B; Table 1), suggesting that the dendritic mislocalization of SNB-1 in *lrk-1* and *apb-3* depends on SYD-2. Unlike the null allele *syd-2*(*ok217*), the loss-of-function allele *syd-2*(*ju37*) does not suppress the dendritic mislocalization of SNB-1 to the ASI dendrite in *lrk-1* and *apb-3* mutants (Fig. 6B). SYD-2 appears to act redundantly with the AP-1 complex to regulate polarized SVp trafficking. Additionally, SYD-2 acts similarly to the AP-1 complex to facilitate SVp entry into dendrites in *lrk-1* and *apb-3* mutants.

We also assessed CTNS-1 localization in dendrites and found that only *lrk-1* shows a significant increase in the number of dendritic CTNS-1 puncta, while *apb-3*, *syd-2*(*ok217*), *unc-101*, and *unc-101; syd-2* are all similar to wildtype (Fig. 6C and 6D). We then assessed whether SYD-2 regulates the trafficking of dendritic cargo, which are known to depend on UNC-101 (Dwyer *et al*., 2001). Both *unc-101* and *apb-3* show mislocalization of the ODR-10::GFP receptor to the AWC axon, while wildtype and *syd-2* mutants localize ODR-10::GFP only along the AWC dendrite and at the dendritic tip (Fig. S4B). We previously showed that UNC-101 regulates the length of the SVp carriers that exit the neuronal cell bodies (Choudhary *et al*., 2017). *unc-101* mutants form longer SVp carriers than that seen in wildtype; however, *syd-2* does not alter the longer SVp carrier length seen in *unc-101* mutants (Fig. S4C).

These data suggest that *syd-2* and *unc-101* are genetically redundant in preventing SVp entry into dendrites but *syd-2* does not alter the axonal phenotypes of *unc-101*. Additionally, the SYD-2 LH1 domain is likely sufficient to enable dendritic entry of atypical SVp carriers formed in *lrk-1* and *apb-3* mutants. Furthermore, *lrk-1* seems to have wider dendritic trafficking defects than those seen in *apb-3*.

### SYD-2 alters the localization of UNC-101 in head neurons

As previously reported, the localization of UNC-101 on the Golgi is altered in *lrk-1* mutants (Choudhary *et al*., 2017) (Fig. 6F). Thus, we examined if *syd-2* alters the localization of UNC-101::GFP in the cell bodies of neurons in the head ganglia (Fig. 6E). The UNC-101::GFP puncta are fainter and smaller (Fig. 6F, 6G and 6H; Suppl. Tables 21 and 22), and a higher percentage of cell bodies have no or fewer puncta in the head neurons in *syd-2* mutants as compared to that in wildtype (Fig. S4D, Suppl. Table 23). In ventral cord neurons, *syd-2* affects the intensity of UNC-101::GFP puncta (Fig. 6I and 6J; Suppl. Table 24). Thus, SYD-2 alters the localization of UNC-101 in *C*. *elegans* neurons, which might account for its role in suppressing the dendritic mistrafficking of some SVps in head neurons.

Further, the loss-of-function allele of *syd-2, syd-2*(*ju37*), did not alter the intensity or size of UNC-101::GFP puncta (Fig. S4E, F, and G; Suppl. Tables 25 and 26). This suggests that the SYD-2 N-terminus domain is sufficient for AP-1 localization.

## Discussion

Our study, like others, shows that SVps are trafficked in many heterogenous carriers and sometimes with lysosomal proteins, suggesting that SVps and lysosomal proteins share trafficking routes (Figs. 1A,1B, 2B, and 2C) (Maeder, Shen and Hoogenraad, 2014; Newell-Litwa *et al*., 2009; Vukoja *et al*., 2018). LRK-1 and, as reported earlier, the AP-3 complex, help in sorting SVps away from lysosomal proteins (Figs. 2B and 2C) (Newell-Litwa *et al*., 2009). In addition, *lrk-1* mutant animals appear to have more widespread trafficking defects of both SVps and lysosomal proteins in comparison to *apb-3* mutants (Fig. 2A and 2F). UNC-104 requires SYD-2 to facilitate the transport of SV carriers that lack lysosomal proteins in wildtype (Fig. 5B) (Zheng *et al*., 2014). The SV-lysosome carrier is dependent on UNC-104, but is largely independent of SYD-2 in wildtype (Fig. S3B). However, in the absence of the AP-3 complex, the preference is switched such that the SV-lysosomes depend on both UNC-104 and SYD-2, but the SVs are only partially dependent on UNC-104 and independent of SYD-2 (Fig. 3B, 3C, 5B, 5C and S3A). Some effects on SYD-2 are likely to be mediated via AP-3 localization to membrane surfaces, perhaps working in concert with UNC-104 to regulate the kinetics of AP-3 membrane cycling. The polarized trafficking of SVps appears to require either SYD-2 or UNC-101, which act redundantly with each other likely due to the role of SYD-2 in enabling localization of the AP-1 complex to the Golgi (Fig. 6B).

LRRK2 is known to affect the trafficking of lysosomal proteins (Kuwahara *et al*., 2016; Piccoli and Volta, 2021; Inoshita *et al*., 2022), SVps (Sakaguchi-Nakashima *et al*., 2007; Cirnaru *et al*., 2014; Choudhary *et al*., 2017), retromer and ER-Golgi proteins (Xiong *et al*., 2012; MacLeod *et al*., 2013; Linhart *et al*., 2014), dense core vesicle proteins (Inoshita *et al*., 2022), RAB GTPases (Steger *et al*., 2016; Lanning *et al*., 2018; Madero-Pérez *et al*., 2018), neurotransmitter transporters (Iovino *et al*., 2022), autophagy-related proteins LC3 and LAMP-1 (Wallings, Connor-Robson and Wade-Martins, 2019), and mitochondria (Weindel *et al*., 2020). The trafficking and localization of lysosomal proteins via LRRK2 seem to depend on RAB-7 and the retromer complex (Dodson *et al*., 2012; Vilariño-Güell *et al*., 2011; Zimprich *et al*., 2011). AP-3 is also known to play a key role in sorting lysosomal proteins in a variety of cells and separating SVps from lysosomal proteins at a common trafficking compartment (Salazar *et al*., 2004; Newell-Litwa *et al*., 2009; Kuwahara *et al*., 2016). Additionally, the *C. elegans* AP-3 complex is shown to be a downstream effector of LRK-1/LRRK2 in axon outgrowth and the co-transport of SNB-1 and RAB-3 along the neuronal process (Kuwahara *et al*., 2016; Choudhary *et al*., 2017). *lrk-1* mutants show more widespread trafficking defects than *apb-3* mutants, such as the presence of LMP-1 along the neuronal process, the presence of CTNS-1 in the dendrite, and reduced cotransport of SNG-1 with RAB-3 (Fig. 2A, 2F and 6C). *lrk-1* and *apb-3* mutants appear to share all other remaining phenotypes, notably that many more SNG-1-transport carriers contain CTNS-1, while nearly all CTNS-1 carriers continue to carry SNG-1 as seen in wildtype (Fig. S2B).

The AP-2 complex is reported to regulate the trafficking of LAMP-1 and LAMP-2 to the lysosomes via the plasma membrane, while the AP-3 complex has little effect on their trafficking (Janvier and Bonifacino, 2005); this supports our data that trafficking of LMP-1 is likely mediated by LRRK2 independently of the AP-3 complex. Thus, LRK-1 may act upstream of AP-3; however, our data do not fully exclude the possibility that LRK-1 and AP-3 act additively to regulate the localization of a subset of lysosomal markers (Fig. S1H-K).

The AP-3 complex can physically bind to LRRK2 (Kuwahara *et al*., 2016; Heaton *et al*., 2020). Therefore, some of the trafficking defects seen in *lrk-1* may occur through its ability to affect the efficient recruitment of the AP-3 complex to membrane surfaces (Fig. 2G-J), as has already been seen for AP-1 (Choudhary *et al*., 2017). Phosphorylation of the AP-3 complex has been shown to be necessary to recruit on SVs and to play a role in endosomal SV biogenesis (Faundez and Kelly, 2000). Further, LRRK2 has been shown to physically interact with the AP-2 complex via its ROC domain (Heaton *et al*., 2020). The LRRK2 ROC domain regulates the LRRK2 kinase activity (Deng *et al*., 2008). Therefore, LRRK2, via its kinase activity (Heaton *et al*., 2020), could regulate AP-3’s localization or activity. Alternatively, LRK-1 could alter the composition of membrane compartments (Piccoli and Volta, 2021) and indirectly affect the recruitment and function of the AP-3 complex.

UNC-104/KIF1A is a critical motor for transporting SVps (Hall and Hedgecock, 1991; Okada and Hirokawa, 1999; Pack-Chung *et al*., 2007). The SV-lysosomes in wildtype, although dependent on UNC-104, do not extend very far into the axon (Fig. 3C, S1H and S3B), perhaps because they have fewer numbers of UNC-104 motors on their surface compared to SVp carriers lacking lysosomal proteins. In *lrk-1* and *apb-3* mutants, SVs partially depend on UNC-104 (Fig. 3B and S3A) (Choudhary *et al*., 2017). It is likely that in *lrk-1* mutants, both SVs and SV-lysosomes depend on multiple motors for their axonal transport, much like that seen in *unc-16* mutants, where UNC-16 acts upstream of LRK-1 (Byrd *et al*., 2001; Brown *et al*., 2009; Choudhary *et al*., 2017). However, *syd-2* mutants, despite sharing some SVp trafficking defects with *lrk-1* and *apb-3* mutants (Fig. 4G and 4H), retain UNC-104 dependence for both SV and SV-lysosome transport (Fig. S3A, 5B and 5C).

SYD-2 is thought to physically associate with the motor, cluster UNC-104, and regulate motor processivity (Shin *et al*., 2003; Wagner *et al*., 2009; Zheng *et al*., 2014; Stucchi *et al*., 2018). The clustering of UNC-104 and increase in processivity might account for the UNC-104 dependence of SVp transport on SYD-2. The effect of SYD-2 on UNC-104-dependent transport may rely on the pre-existing numbers of UNC-104 on the cargo surface. A larger number of motors on the cargo surface may be more sensitive to the UNC-104-clustering activity of SYD-2. Active zone proteins like Piccolo and Bassoon have been thought to cluster vesicles, although some studies suggest that such active zone proteins can be transported in carriers along with SVps (Jin and Garner, 2008; Goldstein, Wang and Schwarz, 2008; Maas *et al*., 2012). SYD-2 is both an UNC-104 interactor and an active zone protein (Zhen and Jin, 1999; Zheng *et al*., 2014). *syd-2* mutants do not show major changes in the localization of SVps or lysosomal proteins and the degree of co-transport of most SV and lysosomal markers assessed (Fig. 4A-D, 4F-H, 5B and S3C-D). This suggests that SYD-2, despite interacting with UNC-104, does not have major roles in the transport or localization of membrane cargo by itself. However, its role is uncovered when there is a reduction in the levels of UNC-104 motor, particularly in the transport of SVs (Fig. 5B and S3A). The reduction in the transport of SV-lysosomes in *apb-3* depends on the presence of an UNC-104-interacting domain of SYD-2 (Fig. 4A, note *apb-3*; *syd-2*(*ju37*)). In the absence of SYD-2’s UNC-104-interacting domain, UNC-104 may not effectively cluster on the surface of SV-lysosomes and therefore, transport of these compartments is reduced. Thus, we think that our data can be explained by SYD-2’s action with UNC-104 rather than a clustering role for multiple vesicles. A role of SYD-2 via regulating the balance/activity of microtubule-dependent motors has also been proposed in lysosome localization in motor neurons of *C. elegans* (Edwards *et al*., 2015b).

Localization of the AP complexes is altered in *syd-2* mutants (Fig. 2G-J, S2G, 6F-H, S4D). There are more and brighter APB-3 puncta in *syd-2*, while there are fewer, less bright, and smaller UNC-101 puncta in *syd-2* animals. The effects of SYD-2 on APB-3 may be explained in two ways. AP-3 recruitment to membrane surfaces depends on binding to cargo proteins (Schoppe *et al*., 2021). Therefore, after AP-3 has sorted cargo, SYD-2 may facilitate UNC-104 clustering, and thereby permit sufficient force generation to enable exit of cargo proteins from an endosomal compartment. Multiple motors are known to generate greater pulling force and deformation of membrane compartments (Roux *et al*., 2002; Du *et al*., 2016). Moreover, the Kinesin 3 family motor KIF13A has been shown to physically bind the AP-1 complex to regulate trafficking of mannose-6-phosphate receptor and the melanosomal cargo, Tyrp1, through affecting AP-1 localization (Nakagawa *et al*., 2000; Delevoye *et al*., 2014). SYD-2’s action may facilitate a similar role of UNC-104 in trafficking. An alternate possibility is that the kinetics of sorting is affected in the absence of SYD-2, leading to persistence of AP-3 complexes on membrane surfaces observed as an increase in the number of puncta in *syd-2* mutants. It is unclear how SYD-2 might influence the recruitment of the AP-1 complex to the membrane. One possibility is that the changes in the AP-3 localization and potential changes in flux through the secretory pathway lead to slowing down of trafficking and therefore changes in localization of AP-1 to reduce cargo jamming in Golgi and post-Golgi compartments.

Polarized trafficking of SVps, specifically their exclusion from dendrites, is dependent on both LRK-1 and the AP-3 complex. SNB-1 mistrafficking in both *lrk-1* and *apb-3* mutants is dependent on SYD-2 as well as the AP-1 complex (Fig. 6B) (Sakaguchi-Nakashima *et al*., 2007; Choudhary *et al*., 2017). The role of SYD-2 in preventing SNB-1 from entering the dendrite in *lrk-1* and *apb-3* mutants might be due to the reduced levels of AP-1 on the Golgi (Fig. 6G, 6H, and 6J). Therefore, in the allele of *syd-*2 that does not affect AP-1 localization, *syd-*2(*ju37*), *lrk-1* and *apb-*3 mutants continue to mistraffick SNB-1 to dendrites (Fig. S4E, F and G). The mistrafficking of SNB-1 into dendrites of *unc-101*; *syd-2* double mutants may be akin to the dendritic mislocalization of SVp in *unc-104* mutants (Yan *et al*., 2013). The lack of sufficient UNC-104 activity may permit dynein motors to enable dendritic entry of SVps.

In conclusion, we propose that in the SV biogenesis pathway, one key step is the separation of SVps from lysosomal proteins via LRK-1 and the AP-3 complex. We also propose a novel role for the active zone protein SYD-2 as a regulator of SVp trafficking, acting downstream to the AP-3 complex and via UNC-104, and as a regulator of polarized distribution of SVps acting along with the AP-1 complex. We show that SYD-2 genetically interacts with and alters the localization of both the AP-3 and AP-1 complexes to regulate the transport and polarized distribution of SVp carriers in *C. elegans* neurons.

## Materials and methods

### Strain maintenance

*C*. *elegans* strains were grown and maintained at 20 °C on NGM plates seeded with *E*. *coli* OP50 strain using standard methods (Brenner, 1974). BD Bacto-Petone and BD Agar for the NGM were sourced from Becton, Dickinson and Company NJ, USA. All Sigma salts and Sigma cholesterol were obtained from local distributors of Sigma and Merck products. L4 or 1-day adult animals were used for imaging in all cases. The strains used are listed in Supplementary Table 1. Some strains were provided by the CGC, which is funded by NIH Office of Research Infrastructure Programs (P40 OD010440).

### Plasmid construction

Expression plasmids were generated using standard PCR-based subcloning techniques. The *mec-4*p::*ctns-1*::*mCherry* plasmid (TTpl 509) was generated by replacing the *unc-129*p from #KG371 (Edwards *et al*., 2013) with *mec-4*p using *Hind*III and *Bam*HI restriction enzymes. The *str-3*p::*ctns-1*::*mCherry* was generated by replacing the *unc-129*p from #KG371 with *str-3*p using *Bam*HI and *Apa*I restriction sites. To generate the *mec-4*p::*sng-1*::*gfp* plasmid (TTpl 696), SNG-1::GFP was amplified from NM491 (Zhao and Nonet, 2001) and cloned into a *mec-4*p containing vector using *Nhe*I and *Eco*RV restriction sites. To generate *rab-3*p::*apb-3*::*gfp* (TTpl 796), APB-3 was amplified from genomic DNA using Phusion Polymerase and cloned into a *rab-3*p-containing vector using *Nhe*I and *Age*I restriction sites. To generate touch neuron specific expression plasmids for *rab-*7 and *lmp-1* under the *mec-7* promoter (Hamelin *et al*., 1992), cloning was performed using the Gateway *in vitro* recombination system (Invitrogen, Carlsbad, CA) using Grant lab modified versions of MiniMos enabled vectors pCFJ1662 (Hygromycin resistant) and pCFJ910 (G418 resistant) (gifts of Erik Jorgensen, University of Utah, Addgene #51482): pCFJ1662 Pmec7 GTWY mNeonGreen let858 (34F6) or pCFJ1662 Pmec7 mNeonGreen GTWY let858 (34D4), and pCFJ910 Pmec7 mScarleti GTWY let858 (33B6). pDONR221 entry vectors containing coding regions for *lmp-1* and *rab-7* were recombined into neuronal destination vectors by Gateway LR clonase II reaction to generate C-/N-terminal fusions. Single-copy integrations were obtained by MiniMOS technology (Frøkjaer-Jensen *et al*., 2008).

### Generation of transgenic *C*. *elegans*

Transgenic lines were generated by following standard microinjection procedure (Fire *et al*., 1998) using an Olympus Ix53 microscope equipped with 20× and 40× lenses, Narishige M-152 micromanipulator (Narishige, Japan), and Eppendorf Femtojet 2 microinjector (local distributors of Eppendorf products). The F2 progeny that inherited and stably expressed the extrachromosomal transgene were UV irradiated to generate integrated lines. Worms showing 100% transmission were selected and outcrossed with the wildtype N2 strain five times. Detailed information on the concentration of plasmids and co-injection markers used is listed in Supplementary Table 2.

### Imaging

**(i) Static imaging:** L4 or 1-day adult worms were immobilized using 30 mM sodium azide and mounted on 2–5% agarose pads. Images were acquired on an Olympus IX73 Epifluorescence microscope with an Andor EMCCD camera or the Olympus Fluoview FV1000 confocal laser scanning microscope or Olympus IX83 with Perkin Elmer Ultraview Spinning Disc confocal microscope fitted with a Hamamatsu EMCCD camera. Since AP-3 localization is sensitive to levels of ATP (Faundez and Kelly, 2000), static imaging of APB-3::GFP was performed using 5 mM Tetramisole. APB-3::GFP was imaged on Olympus Spin SR10 (SoRA, 50 μm disk) fitted with Teledyne Photometrics sCMOS camera.
**(ii) Time-lapse imaging:** L4 worms were anesthetized in 3 mM tetramisole (Sigma-Aldrich) and mounted on 5% agarose pads. Time-lapse images were acquired in Olympus IX83 with Perkin Elmer Ultraview Spinning Disc confocal microscope and a Hamamatsu EMCCD camera or the Olympus Fluoview FV1000 confocal laser scanning microscope. Dual color simultaneous imaging was performed at 3 frames per second (fps), dual color sequential imaging was done at 1.3 fps, and single fluorophore imaging for analysis of vesicle length was done at 5 fps. All movies were 3 minutes long, and the region of imaging in the PLM comprised the first 60–100 μm of the neuronal process immediately outside the cell body, with the cell body in the frame of imaging. Live imaging of EBP-2::GFP to assess microtubule polarity was carried out using an Olympus IX73 Epifluorescence microscope with an Andor EMCCD camera at 3 fps.

### Analysis

All analysis was done using FIJI (Schindelin *et al*., 2012).

**(i) Co-migration analysis:** Kymographs were generated from identical regions of the movie in both color channels utilizing the ImageJ plugin MultipleKymograph. The kymographs were then synchronized and the overlapping sloped lines were considered as co-migrating particles. For dual-color co-migration analysis, number of moving vesicles were counted which were positive for GFP alone, RFP alone, and vesicles positive for both GFP and RFP. Total number of vesicles = number of vesicles positive only for GFP + number of vesicles only positive for RFP + number of vesicles positive for both GFP and RFP.

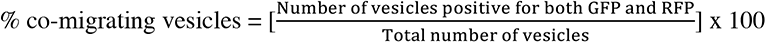

Fraction of GFP-positive vesicles co-migrating with RFP-positive vesicles =

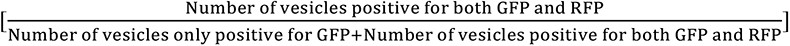

For detailed methods, please refer to (Nadiminti and Koushika, 2022).

**(iii) Quantitation of penetrance of CTNS-1 puncta that exit into PLM neurites:** For each genotype, at least 30 animals were annotated to observe the extent of CTNS-1 (or RAB-7 or LMP-1) presence in the PLM major neurite. Penetrance was measured by calculating the number of animals in which CTNS-1 (or RAB-7 or LMP-1) was present at or beyond the first 25 μm and 50 μm away from the cell body.
**(iv) Quantitating the direction of motion of CTNS-1-carrying compartments:** Only moving CTNS-1-carrying compartments were analyzed for their direction of motion. For CTNS-1-marked compartments moving clearly in a particular direction, they were annotated as such. For those moving bidirectionally, their net displacement was used to identify their direction of motion. If the vesicle’s final position at the end of the kymograph was closer to the cell body than when it started, it was considered to have moved retrogradely. If the vesicle’s final position at the end of the kymograph was farther away from the cell body than when it started, it was considered to have moved anterogradely. For vesicles whose position at the end of the kymograph remained largely unchanged, they were either not considered for analysis or were assigned the direction in which they were moving immediately before the end of the kymograph, depending upon how discernible their direction of motion was.
**(iv) Density of CTNS-1 in the ASI dendrite:** The number of CTNS-1 puncta in the dendrite and the length of measurable region (ROI) in the dendrite from the cell body to the end was counted for each animal. The density of lysosomes per 10 μm was calculated as:

[Number of CTNS-1 puncta in the dendrite/Length of the dendrite ROI] × 10

**(iv) Quantitation of intensity of UNC-101::GFP and APB-3::GFP puncta:** For UNC-101::GFP, two regions were chosen – (i) the cell bodies of the head neuron ganglia and (ii) the cell bodies along the ventral nerve cord. For APB-3::GFP, neurons in three regions – the head, along the ventral cord, and the tail – were analyzed. For both UNC-101::GFP and APB-3::GFP, per cell body, the number of puncta was calculated on a plane with the best focus for that cell body. On the same plane, the size and intensity of each puncta were measured. A cytosolic region close to one of the puncta was chosen to measure puncta/cytosolic intensity. Puncta intensity was quantitated by dividing the intensity of each puncta by the cytosolic intensity. All the values of puncta intensity to cytosolic intensity per cell body were averaged and plotted.
**(v) Vesicle length analysis:** In every kymograph, random non-overlapping ROIs (regions of interest) were chosen to measure the size of the vesicles. These random ROIs were generated by [Macro 1]. Any macro-generated random ROI that overlapped with a previous ROI for that kymograph was not used for the analysis. Within each ROI, the length of each moving compartment was quantified by measuring the thickness of the sloped line along the x-axis. Such measurements were done at regions not overlapping with stationary particles or other moving particles.
**(vi) Microtubule polarity:** Kymographs were generated from live movies of EBP-2::GFP in the axonal and anterior dendritic regions of the PVD neuron imaged at 3 fps. The number of anterogradely and retrogradely moving EBP-2 were counted from the kymographs and plotted.

### Statistical analysis

All statistical analyses were performed using OriginLab 2019. Distributions were checked for normality using the Shapiro–Wilk test. Data that fit a normal distribution were compared using one-way ANOVA with Tukey’s post-hoc test. Data that did not fit a normal distribution were compared using the Mann–Whitney test. Differences were considered significant when the p-value < 0.05.

## Supporting information

All Supplemental tables

## Acknowledgments

We thank Dr. Kenneth Miller for the CTNS-1 plasmid, Dr. Hidenori Taru for the SYD-2 deletion strains and constructs, and Drs. Mei Zheng and Yishi Jin. We thank Dr. Michael Nonet for SYD-2 constructs and the *mec*-*7*p::*snb-1*::*gfp* plasmid. We thank Badal Singh Chauhan for generating the transgenic strain *tbEx384* [*mec*-*7*p::*snb-1*::*gfp*]. Some strains were provided by the CGC, which is funded by NIH Office of Research Infrastructure Programs (P40 OD010440). Research in the SPK lab is supported by grants from DAE (1303/2/2019/R&D-II/DAE/2079) and PRISM (12-R&D-IMS-5.02-0202). Research in the BDG lab is supported by the NIH grant R01GM135326.

## List of supplementary movies and legends

**Supplementary movie 1: CTNS-1 and SNG-1 in WT**

SNG-1::GFP and CTNS-1::mCherry in the PLM neuronal process. Imaged sequentially at 1.3 frames per second (fps), playback at 20 fps. Genotype: wildtype. Cell body on the right.

**Supplementary Movie 2: RAB-7 and SNG-1 in WT**

SNG-1::GFP and mScarlet::RAB-7 in the PLM neuronal process. Imaged sequentially at 1.3 frames per second (fps), playback at 20 fps. Genotype: wildtype. Cell body on the right.

**Supplementary Movie 3: CTNS-1 and SNG-1 in *lrk-1***

SNG-1::GFP and CTNS-1::mCherry in the PLM neuronal process. Imaged sequentially at 1.3 frames per second (fps), playback at 20 fps. Genotype: *lrk-1*(*km17*). Cell body on the right.

**Supplementary Movie 4: CTNS-1 and SNG-1 in *apb-3***

SNG-1::GFP and CTNS-1::mCherry in the PLM neuronal process. Imaged sequentially at 1.3 frames per second (fps), playback at 20 fps. Genotype: *apb-3*(*ok429*). Cell body on the right.

**Supplementary Movie 5: RAB-7 and SNG-1 in *lrk-1***

SNG-1::GFP and mScarlet::RAB-7 in the PLM neuronal process. Imaged sequentially at 1.3 frames per second (fps), playback at 20 fps. Genotype: *lrk-1*(*km17*). Cell body on the right.

**Supplementary Movie 6: CTNS-1 and SNG-1 in *unc-104***

SNG-1::GFP and CTNS-1::mCherry in the PLM neuronal process. Imaged sequentially at 1.3 frames per second (fps), playback at 20 fps. Genotype: *unc-104*(*e1265tb120*). Cell body on the right.

**Supplementary Movie 7: CTNS-1 and SNG-1 in *syd-2***

SNG-1::GFP and CTNS-1::mCherry in the PLM neuronal process. Imaged sequentially at 1.3 frames per second (fps), playback at 20 fps. Genotype: *syd-2*(*ok217*). Cell body on the right.

---

**Supplementary Figure 1:**
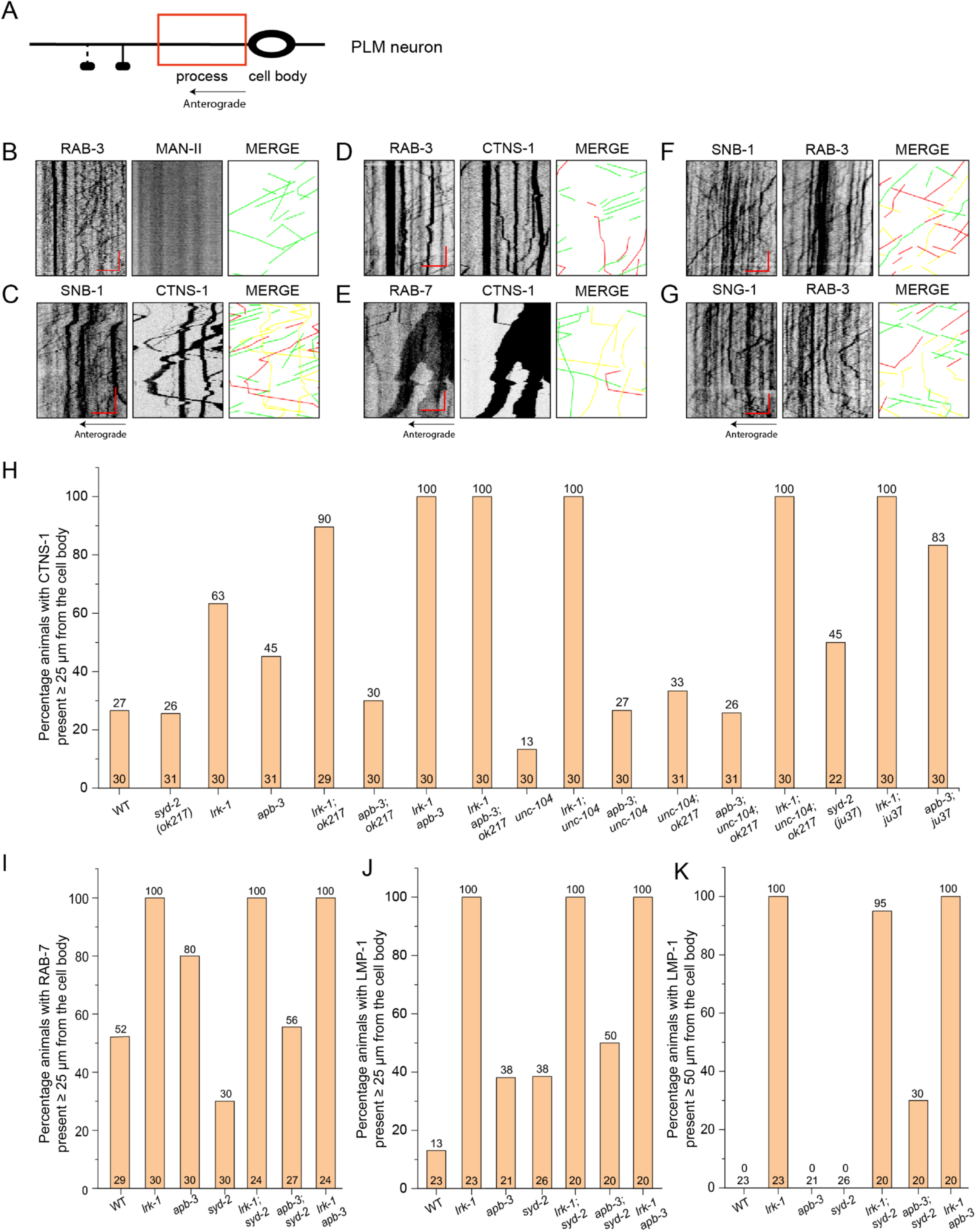
(A) Schematic of the PLM neuron. The red box highlights the region of imaging in the proximal major neuronal process. The arrow indicates the direction of anterograde motion, away from the cell body into the neuronal process. (B) Kymographs from dual-color imaging of RAB-3 with MAN-II in WT, imaged simultaneously at 3 frames per second (fps). Green traces indicate moving RAB-3 vesicles. Scale bars x-axis: 5 μm, y-axis: 30 s. (C) Kymographs from dual-color imaging of SNB-1 with CTNS-1 in WT, imaged sequentially at 1.3 fps. Green traces indicate moving SNB-1 vesicles, yellow traces indicate moving vesicles co-transporting SNB-1 and CTNS-1, and red traces indicate moving CTNS-1 vesicles. Scale bars x-axis: 5 μm, y-axis: 30 s. (D) Kymographs from dual-color imaging of RAB-3 with CTNS-1, imaged simultaneously at 3 fps. Green traces indicate moving RAB-3 vesicles, yellow traces indicate moving vesicles co-transporting RAB-3 and CTNS-1, and red traces indicate moving CTNS-1 vesicles. Scale bars x-axis: 5 μm, y-axis: 10 s. (E) Kymographs from dual-color imaging of mNeonGreen::RAB-7 with CTNS-1::mCherry, imaged sequentially at 1.3 fps. Green traces indicate moving RAB-7 vesicles, yellow traces indicate moving vesicles co-transporting RAB-7 and CTNS-1, and red traces indicate moving CTNS-1 vesicles. Scale bars x-axis: 5 μm, y-axis: 30 s. (F) Kymographs from dual-color imaging of SNB-1 with RAB-3, imaged simultaneously at 3 fps. Green traces indicate moving SNB-1 vesicles, yellow traces indicate moving vesicles co-transporting SNB-1 and RAB-3, and red traces indicate moving RAB-3 vesicles. Scale bars x-axis: 5 μm, y-axis: 10 s. (G) Kymographs from dual-color imaging of SNG-1 with RAB-3, imaged sequentially at 1.3 fps. Green traces indicate moving SNG-1 vesicles, yellow traces indicate moving vesicles co-transporting SNG-1 and RAB-3, and red traces indicate moving RAB-3 vesicles. Scale bars x-axis: 5 μm, y-axis: 30 s. (H) Penetrance for the number of animals in which CTNS-1 localizes up to 25 μm of the PLM neuronal process away from the cell body. Numbers inside the bars indicate the number of animals per genotype. Numbers above the bars indicate the penetrance values. For bar graphs with very little height, the lower number indicates the number of animals for that genotype while the number above indicates the penetrance value. (I) Penetrance for the number of animals in which RAB-7 localizes up to 25 μm of the PLM neuronal process away from the cell body. Numbers inside the bars indicate the number of animals per genotype. Numbers above the bars indicate the penetrance values. (J) and (K) Penetrance for the number of animals in which LMP-1 localizes up to 25 μm and 50 μm, respectively, of the PLM neuronal process away from the cell body. Numbers inside the bars indicate the number of animals per genotype. Numbers above the bars indicate the penetrance values. For bar graphs with very little height, the lower number indicates the number of animals for that genotype while the number above indicates the penetrance value.

**Supplementary Figure 2:**
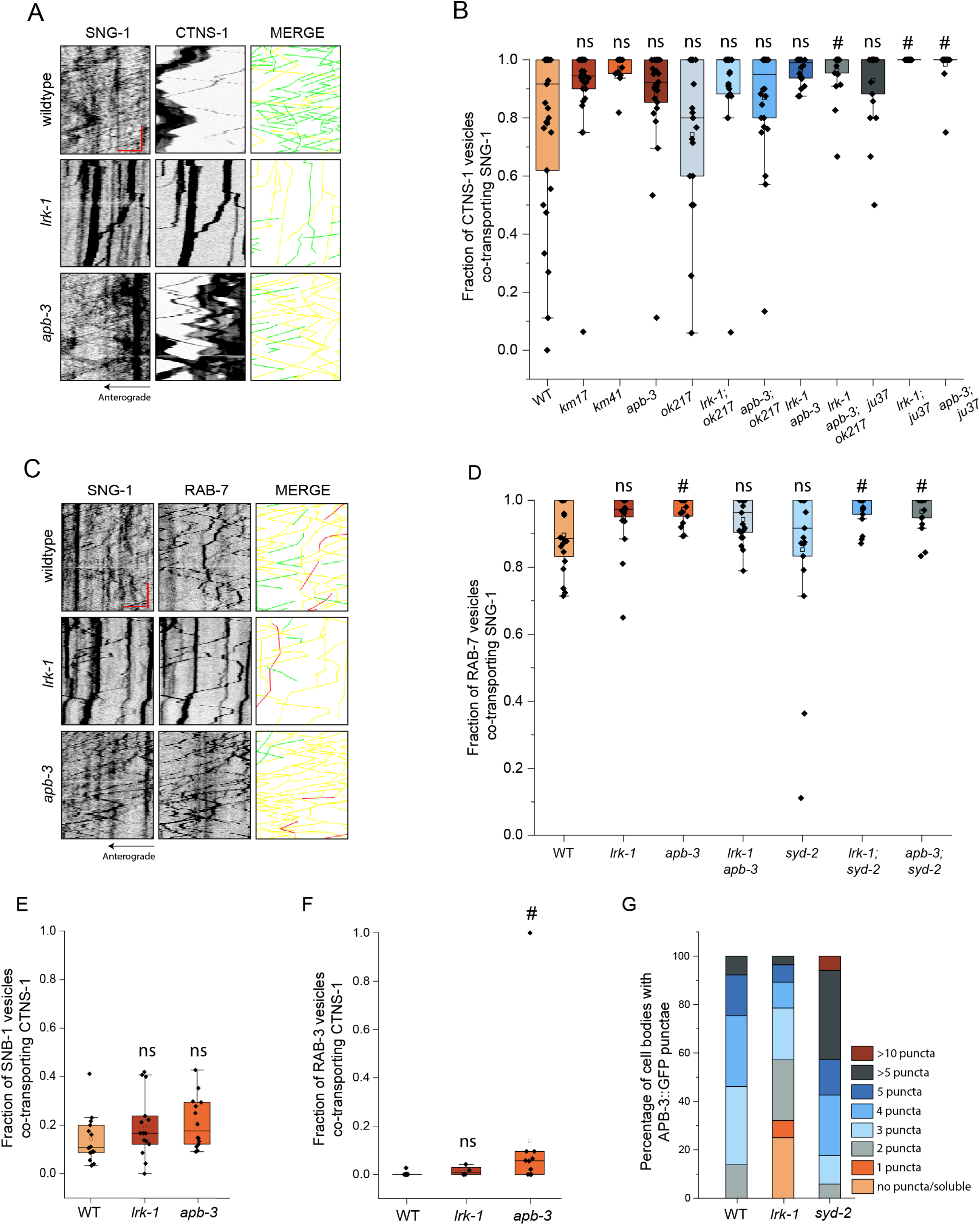
(A) Kymographs from sequential dual-color imaging of SNG-1 and CTNS-1 at 1.3 fps in WT, *lrk-1*(*km17*), and *apb-3*(*ok429*). Green traces indicate moving SNG-1-carrying vesicles, yellow traces indicate moving vesicles co-transporting SNG-1 and CTNS-1, and red traces indicate moving CTNS-1-carrying vesicles. Scale bar x-axis: 5 μm and y-axis: 30 s. (B) Quantitation of fraction of CTNS-1 co-transporting SNG-1 from kymograph analysis of dual color imaging. ^#^P-values ≤ 0.05 (Mann–Whitney Test, all comparisons to WT); ns: not significant; Number of animals per genotype (N) ≥ 20; Number of vesicles (n) > 400. (C) Kymographs from sequential dual-color imaging of SNG-1 and RAB-7 at 1.3 fps in WT, *lrk-1*(*km17*), and *apb-3*(*ok429*). Green traces indicate moving SNG-1-carrying vesicles, yellow traces indicate moving vesicles co-transporting SNG-1 and RAB-7, and red traces indicate moving RAB-7-carrying vesicles. Scale bar x-axis: 5 μm and y-axis: 30 s. (D) Quantitation of fraction of RAB-7 co-transporting SNG-1 from kymograph analysis of dual color imaging. # P-values ≤ 0.05 (Mann–Whitney Test, all comparisons to WT); ns: not significant; N ≥ 20 per genotype; n > 400. (E) Quantitation of co-transport of SNB-1 and CTNS-1 in WT, *lrk-1*(*km17*), and *apb-3*(*ok429*) from kymograph analysis of dual color imaging. P-values > 0.05 (One-Way ANOVA with Tukey’s post-hoc test, all comparisons to WT); ns: not significant; N ≥ 15 per genotype; n > 400. (F) Quantitation of co-transport of RAB-3 and CTNS-1 in WT, *lrk-1*(*km17*), and *apb-3*(*ok429*) from kymograph analysis of simultaneous dual color imaging at 3 fps. # P-values ≤ 0.05 (Mann–Whitney Test, all comparisons to WT); ns: not significant; N = 5 per genotype; n > 500. (G) Percentages of cell bodies of WT, *lrk-1,* and *syd-2* with APB-3::GFP puncta. N > 10 per genotype; n > 75 cell bodies.

**Supplementary Figure 3:**
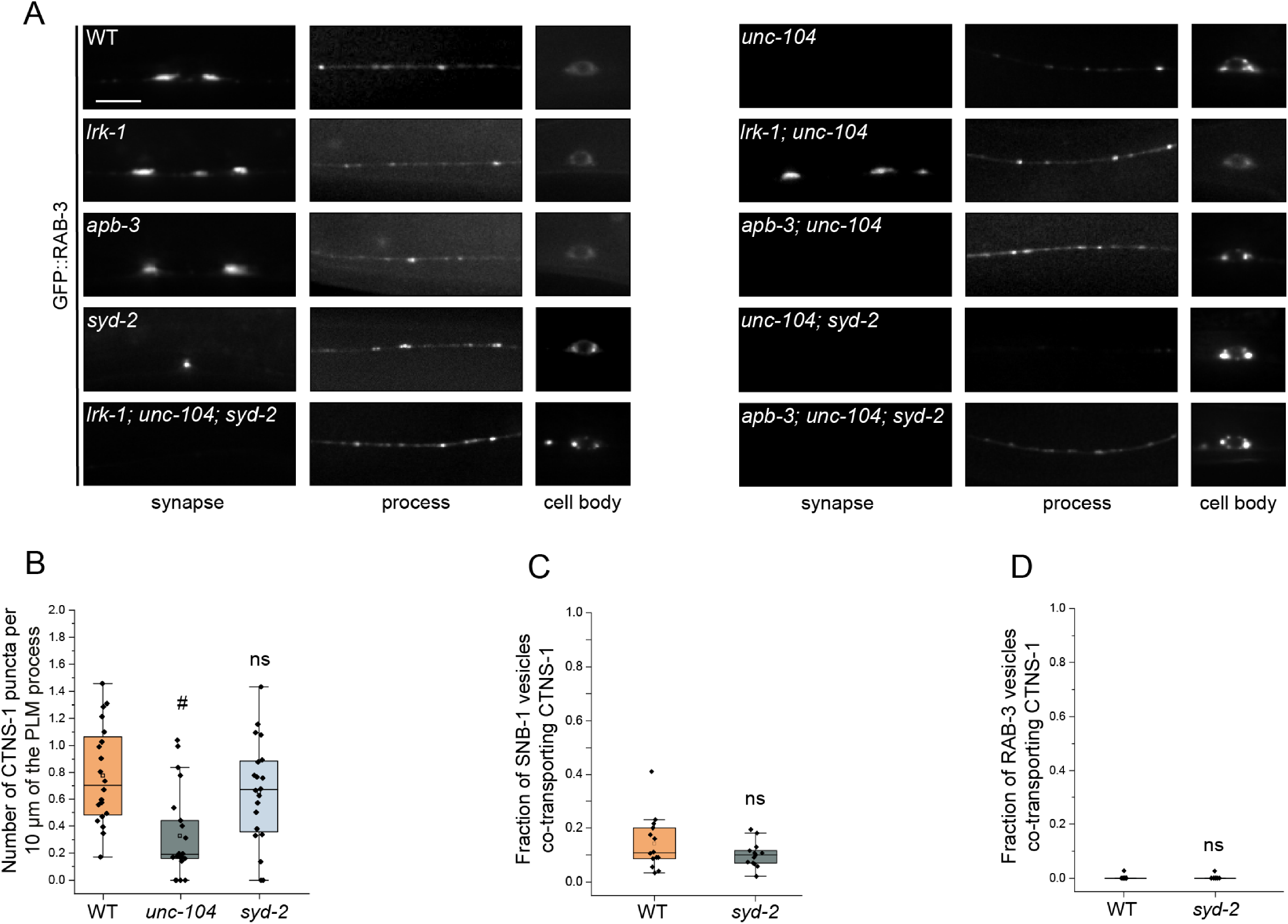
(A) GFP::RAB-3 in the cell body, process, and synapses of PLM neurons showing dependence on UNC-104 in *lrk-1*(*km17*), *apb-3*(*ok429*), and *syd-2*(*ok217*)mutants, and their doubles with *unc-104*(*e1265tb120*). Scale bar: 10 μm. (B) Quantitation of the number of CTNS-1-labelled compartments per 10 μm of the PLM major neurite proximal to the cell body in WT, *unc-104*(*e1265tb120*), and *syd-2*(*ok217*). # P-values ≤ 0.05 (Mann–Whitney Test, all comparisons to WT); ns: not significant; Number of animals (N) ≥ 20 per genotype; Number of CTNS-1-labelled compartments (n) ≥ 70. (C) Quantitation of co-transport of SNB-1 and CTNS-1 in WT and *syd-2*(*ok217*), from kymograph analysis of sequential dual color imaging at 1.3 fps. P-value > 0.05 (Mann– Whitney Test); ns: not significant; N > 15 per genotype; n > 750 vesicles. (D) Quantitation of co-transport of RAB-3 and CTNS-1, in WT and *syd-2*(*ok217*), from kymograph analysis of simultaneous dual color imaging at 3 fps. P-value > 0.05 (Mann– Whitney Test); ns: not significant; N = 5; n > 500 vesicles.

**Supplementary Figure 4:**
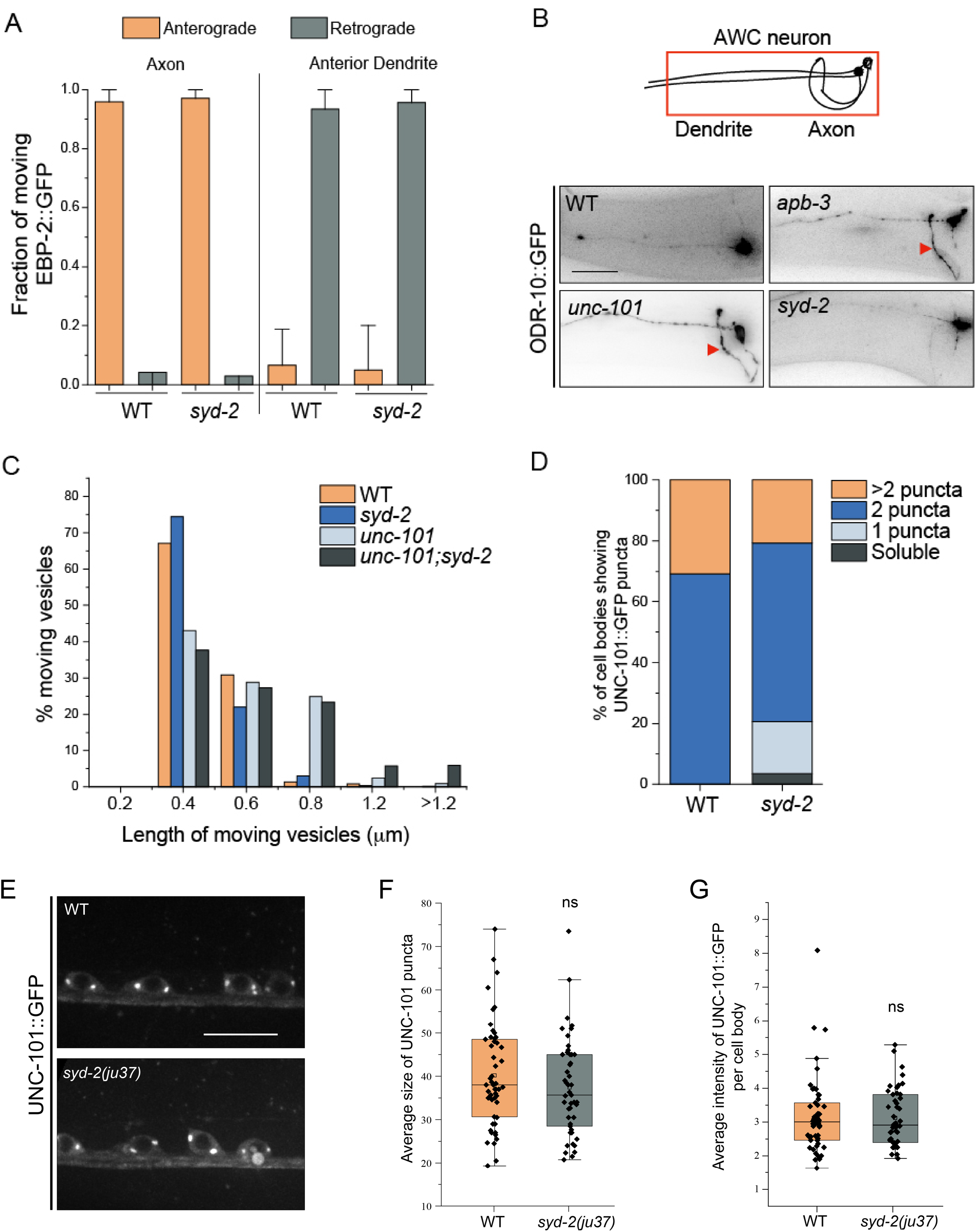
(A) Quantitation of fraction of EBP-2::GFP comets moving in either anterograde or retrograde directions in both the axon and the anterior dendrite of WT and *syd-2*(*ok217*); Number of animals (N) > 8 for each genotype; Number of comets analyzed (n) > 150. (B) ODR-1::GFP in the dendrite and axon of the AWC neuron. Red arrow points to the ODR-1::GFP signal in the AWC axon in *syd-2*(*ok217*), *apb-3*(*ok429*), and *unc-101*(*m1*). Scale bar: 20 μm. (C) Quantitation of sizes of moving RAB-3 containing SVp carriers in WT, *syd-2*(*ok217*), *unc-101*(*m1*), and *unc-101*; *syd-2*. The x-axis depicts the length (in μm) of moving RAB-3 carrying SVp carriers. The y-axis depicts the percentage of moving RAB-3 carrying SVp carriers of various lengths. Number of animals (N) ≥ 9 per genotype; Number of vesicles (n) > 400. (D) Quantitation of the number of UNC-101::GFP puncta per cell body in WT and *syd-2*(*ok217*). P-value > 0.05 (Mann–Whitney Test); N > 5 animals; n > 25 cell bodies. (E) Images showing UNC-101::GFP puncta in the cell bodies of the ventral nerve cord neurons in WT and *syd-2*(*ju37*). Scale bar: 10 μm. (F) Quantitation of the average size of UNC-101::GFP puncta per cell body in WT and *syd-2*(*ju37*). P-value < 0.05 (Mann–Whitney Test); ns: not significant; N > 5 animals; n > 25 cell bodies. (G) Quantitation of intensity of UNC-101::GFP puncta in the cell bodies of the ventral nerve cord in WT and *syd-2*(*ju37*). The ratio of the intensity of UNC-101::GFP puncta to cytosolic intensity in the cell body is plotted. P-value < 0.05 (Mann–Whitney test); ns: not significant; N > 5 animals; n > 10 cell bodies.

