## Supplementary material for "Active zone protein SYD-2/Liprin-α acts downstream of LRK-1/LRRK2 to regulate polarized trafficking of synaptic vesicle precursors through clathrin adaptor protein complexes": All Supplemental tables

**Supplementary Table 1: List of strains**

| S.No | Strain number | Genotype | Reference |
|---|---|---|---|
| 1 | N2 | Bristol wild type | (Brenner, 1974) |
| 2 | NM2689 | *jsIs821* [*mec-7*p::*gfp*::*rab-3*] | (Bounoutas *et al*., 2009) |
| 3 | TT1555 | *tbEx254* [*mec-7*p::*aman-II*::*mCherry*]; *jsIs821* | (Choudhary *et al*., 2017) |
| 4 | TT2884 | *tbIs388* [*mec-4*p::*sng-1*::*egfp*] | This study |
| 5 | TT2903 | *tbIs381* [*mec-4*p::*ctns-1*::*mCherry*] | This study |
| 6 | TT3029 | *pwSi113* [*mec-7*p::*mScarlet*::*rab-7*::*let858*] G418R | Barth Grant, this study |
| 7 | TT3007 | *tbIs414* [*mec-7*p::*snb-1*::*egfp*] | This study |
| 8 | TT1884 | *tbIs227* [*mec-4*p::*mCherry*::*rab-3*] | (Choudhary *et al*., 2017) |
| 9 | NM664 | *jsIs37* [*mec-7*p::*snb-1*::*gfp*] | (Nonet, 1999) |
| 10 | TT2710 | *tbEx364* [*mec-4*p::*sng-1*::*egfp*] | This study |
| 11 | TT1519 | *tbEx272* | This study |
| 12 | TT2918 | *tbEx384* | This study |
| 13 | TT3286 | *rab-7::mNeonGreen; tbIs381* | Barth Grant, this study |
| 14 | TT3035 | *pwSi225* [*mec-7*p::*lmp-1*::*mNeonGreen*::*let858*]; hygR | Barth Grant, this study |
| 15 | NM2156 | *kyIs105* (str-3p::SNB-1::GFP) | This study |
| 16 | TT2636 | *tbEx316* [*str-3*p::*ctns-1*::*mCherry*] | This study |
| 17 | TT1893 | *oqEx* [*unc-101*p::*unc-101*::*gfp*+pRF4] | (Kaplan *et al*., 2010) |
| 18 | TT2052 | *kyEx3233* [*des2*p::*ebp-2*::*gfp*] | (Maniar *et al*., 2012) |
| 19 | TT1612 | *kyIs156* [*str-1*p::*odr-10*::*gfp*] | (Dwyer *et al*., 2001) |
| 20 | TT3371 | *tbEx486* [*rab-3*p::*apb-3*::*gfp*] | This study |
| 21 | TT1218 | *lrk-1*(*km17*) | (Sakaguchi-Nakashima *et al*., 2007) |
| 22 | TT1225 | *lrk-1*(*km41*) | |
| 23 | TT2060 | *apb-3*(*ok429*) | (Consortium, 2012) |
| 24 | TT1134 | *syd-2*(*ok217*) | (Consortium, 2012) |
| 25 | TT148 | *syd-2*(*ju37*) | (Zhen and Jin, 1999) |
| 26 | TT385 | *unc-104*(*e1265tb120*) | (Kumar *et al*., 2010) |
| 27 | TT1211 | *unc-101*(*m1*) | (Lee, Jongeward and Sternberg, 1994) |

**Supplementary Table 2: List of plasmids**

| | Strain name | Details | Plasmid | Plasmid details | Source |
|---|---|---|---|---|---|
| 1 | *tbEx272* | Generated by injecting TTpl509 (5 ng/µL) with *myo-2*p::*gfp* (50 ng/µL) as coinjection marker. | TTpl509 | *mec-4*p::*ctns-1*::*mCherry*. *unc-129*p in KG#371 was replaced with *mec-4*p using HindIII and BamHI restriction sites. | This study. (Edwards *et al*., 2013) KG#371 was a gift from K G Miller. |
| 2 | *tbEx369* | Generated by injecting TTpl509 (20 ng/µL) with *myo-2*p::*gfp*::*h2b* (TTpl592) (40 ng/µL) as coinjection marker. | | | |
| 3 | *tbEx364* | Generated by injecting TTpl696 (5 ng/µL) with *myo-2*p::*mCherry* (TTpl580) (10 ng/µL) as coinjection marker. | TTpl696 | *mec-4*p::*sng-1*::*gfp*. SNG-1::GFP was amplified from NM491 and cloned into a *mec-4*p containing vector using NheI and EcoRV restriction sites. | This study. NM491 [pSY3 (SNG-1::GFP construct)]. (Zhao and Nonet, 2001) NM491 was a gift from Michael Nonet. |
| 4 | *tbEx316* | Generated by injecting TTpl653 (20 ng/µL) with *unc-122*p::*gfp* (TTpl572) (30 ng/µL) as coinjection marker. | TTpl635 | *str-3*p::*ctns-1*::*mCherry*. *mec-4*p from TTpl509 was switched out for *str-3*p using BamHI and ApaI restriction sites. | This study. TTpl572 was a gift from Kavita Babu. |
| 5 | *tbEx486* | Generated by injecting TTpl796 (Isolate 2) (30 ng/µL) with *myo-2*p::*mCherry* (TTpl580) (10 ng/µL) as coinjection marker. | TTpl796 | *rab-3*p::*apb-3*::*gfp*. APB-3 was amplified from genomic DNA using Phusion Polymerase and cloned into a *rab-3*p containing vector (TTpl698) using NheI and AgeI restriction sites. | This study. |
| 6 | *tbEx384* | Generated by injecting TTpl684 (10 ng/µL) with *myo-2*p::*mCherry* (TTpl580) (10 ng/µL) as coinjection marker. | TTpl684 | *mec-4*p::*snb-1*::*gfp* | This study. Plasmid was a gift from Michael Nonet. |

**Supplementary Table 3: (Associated with Fig. 2A)**

| Fraction of SNG-1 co-migrating with RAB-3 | | |
|---|---|---|
| **Genotype** | **P-value** | **Significant** |
| **One Way ANOVA with Tukey's post Hoc test** | | |
| WT vs *lrk-1* | 0.04 | Yes |
| WT vs *apb-3* | 0.13 | No |
| *lrk-1* vs *apb-3* | 1.4E-05 | Yes |

**Supplementary Table 4: (Associated with Fig. 2B)**

| Fraction of SNG-1 co-migrating with CTNS-1 | | |
|---|---|---|
| **Genotype** | **P-value** | **Significant** |
| **Mann-Whitney Test** | | |
| WT vs *lrk-1(km17)* | 3.2E-08 | Yes |
| WT vs *apb-3(ok429)* | 9.2E-09 | Yes |
| WT vs *lrk-1*(*km17*) *apb-3* | 1.4E-07 | Yes |
| *lrk-1(km17)* vs *lrk-1 apb-3* | 0.26 | No |
| WT vs *lrk-1*(*km41*) | 2.9E-07 | Yes |
| *lrk-1*(*km17*) vs *lrk-1*(*km41*) | 0.008 | Yes |
| **One Way ANOVA with Tukey's post Hoc test** | | |
| *apb-3* vs *lrk-1*(*km17*) *apb-3* | 0.062 | No |
| *lrk-1*(*km41*) vs *apb-3* | 0.58 | No |

**Supplementary Table 5: (Associated with Fig. S2B)**

| Fraction of CTNS-1 co-migrating with SNG-1 | | |
|---|---|---|
| **Genotype** | **P-value** | **Significant** |
| **Mann-Whitney Test** | | |
| WT vs *lrk-1(km17)* | 0.33 | No |
| WT vs *lrk-1(km41)* | 0.06 | No |
| WT vs *apb-3(ok429)* | 0.60 | No |
| WT vs *lrk-1 apb-3* | 0.07 | No |
| *lrk-1(km17)* vs *lrk-1 apb-3* | 0.15 | No |
| *lrk-1(km17)* vs *lrk-1 apb-3; syd-2* | 0.03 | Yes |
| *apb-3* vs *apb-3; syd-2(ok217)* | 0.86 | No |
| WT vs *syd-2(ju37)* | 0.063 | No |
| WT vs *apb-3; syd-2(ju37)* | 0.0056 | Yes |
| WT vs *lrk-1; syd-2(ju37)* | 0.0016 | Yes |
| *syd-2(ju37)* vs *apb-3; syd-2(ju37)* | 0.23 | No |
| *syd-2(ju37)* vs *lrk-1; syd-2(ju37)* | 0.09 | No |
| WT vs *unc-104(e1265tb120)* | 0.23 | No |
| WT vs *lrk-1; unc-104(e1265tb120)* | 0.09 | No |
| WT vs *apb-3; unc-104(e1265tb120)* | 0.03 | Yes |
| WT vs *unc-104(e1265tb120); syd-2(ok217)* | 0.03 | Yes |
| *lrk-1(km17)* vs *lrk-1; unc-104(e1265tb120)* | 0.3 | No |
| *apb-3(ok429)* vs *apb-3; unc-104(e1265tb120)* | 0.006 | Yes |
| *syd-2(ok217)* vs *unc-104(e1265tb120); syd-2* | 0.006 | Yes |
| *unc-104(e1265tb120)* vs *unc-104; syd-2(ok217)* | 0.46 | No |
| *unc-104(e1265tb120)* vs *lrk-1*; *unc-104* | 0.96 | No |
| *unc-104(e1265tb120)* vs *apb-3; unc-104* | 0.42 | No |
| *lrk-1(km17)* vs *lrk-1; syd-2(ju37)* | 0.0002 | Yes |
| *apb-3; syd-2(ok217)* vs *apb-3; syd-2(ju37)* | 0.02 | Yes |
| *syd-2(ju37)* vs *unc-104(e1265tb120)* | 0.78 | No |
| *syd-2(ok217)* vs *syd-2(ju37)* | 0.013 | Yes |
| *apb-3* vs *lrk-1 apb-3* | 0.04 | Yes |
| *apb-3* vs *lrk-1 apb-3; syd-2* | 0.01 | Yes |
| *apb-3* vs *apb-3; syd-2(ju37)* | 0.00068 | Yes |

| | | |
|---|---|---|
| *lrk-1; syd-2(ok217)* vs *lrk-1; syd-2(ju37)* | 0.01 | Yes |

**Supplementary Table 6: (Associated with Fig. 2C)**

| Fraction of SNG-1 co-migrating with RAB-7 | | |
|---|---|---|
| **Genotype** | **P-value** | **Significant** |
| **Mann-Whitney Test** | | |
| WT vs *lrk-1 apb-3* | 5.3E-08 | Yes |
| *lrk-1* vs *lrk-1 apb-3* | 0.0016 | Yes |
| *apb-3* vs *lrk-1 apb-3* | 1.8E-06 | Yes |
| **One Way ANOVA with Tukey's post Hoc test** | | |
| WT vs *lrk-1(km17)* | 0 | Yes |
| WT vs *apb-3(ok429)* | 1.5E-06 | Yes |
| *apb-3* vs *lrk-1* | 0.019 | Yes |

**Supplementary Table 7: (Associated with Fig. S2D)**

| Fraction of RAB-7 co-migrating with SNG-1 | | |
|---|---|---|
| **Genotype** | **P-value** | **Significant** |
| **Mann-Whitney Test** | | |
| WT vs *lrk-1* | 0.11 | No |
| WT vs *apb-3* | 0.02 | Yes |
| WT vs *lrk-1 apb-3* | 0.14 | No |
| WT vs *syd-2* | 0.83 | No |
| WT vs *lrk-1; syd-2* | 0.02 | Yes |
| WT vs *apb-3; syd-2* | 0.05 | Yes |
| *lrk-1* vs *lrk-1; syd-2* | 0.31 | No |
| *lrk-1* vs *lrk-1 apb-3* | 0.57 | No |
| *apb-3* vs *apb-3; syd-2* | 0.98 | No |
| *apb-3* vs *lrk-1 apb-3* | 0.19 | No |
| *syd-2* vs *lrk-1; syd-2* | 0.07 | No |
| *syd-2* vs *apb-3; syd-2* | 0.13 | No |
| *lrk-1* vs *apb-3* | 0.56 | No |
| *lrk-1* vs *syd-2* | 0.26 | No |
| *apb-3* vs *syd-2* | 0.08 | No |

**Supplementary Table 8: (Associated with Fig. S2E)**

| Fraction of SNB-1 co-migrating with CTNS-1 | | |
|---|---|---|
| **Genotype** | **P-value** | **Significant** |
| **Mann-Whitney Test** | | |
| WT vs *lrk-1* | 0.25 | No |
| WT vs *apb-3* | 0.053 | No |
| *lrk-1* vs *apb-3* | 0.9 | No |

**Supplementary Table 9: (Associated with Fig. S2F)**

| Fraction of RAB-3 co-migrating with CTNS-1 | | |
|---|---|---|
| **Genotype** | **P-value** | **Significant** |
| **Mann-Whitney Test** | | |
| WT vs *lrk-1* | 0.28 | No |
| WT vs *apb-3* | 0.036 | Yes |
| *lrk-1* vs *apb-3* | 0.11 | No |

**Supplementary Table 10: (Associated with Fig. 2H)**

| Analyses of APB-3::GFP puncta - number per cell body, size and intensity | | | |
|---|---|---|---|
| **Analysis** | **Genotype** | **P-value** | **Significant** |
| | **Mann-Whitney Test** | | |
| Number of puncta per cell body | WT vs *lrk-1* | 0.00008 | Yes |
| | WT vs *syd-2* | 0.00001 | Yes |
| | *lrk-1* vs *syd-2* | 0.00001 | Yes |

**Supplementary Table 11: (Associated with Fig. 2I)**

| Analyses of APB-3::GFP puncta - number per cell body, size and intensity | | | |
|---|---|---|---|
| **Analysis** | **Genotype** | **P-value** | **Significant** |
| | **Mann-Whitney Test** | | |
| Average size of puncta | WT vs *lrk-1* | 0.00252 | Yes |
| | WT vs *syd-2* | 0.00386 | Yes |
| | *lrk-1* vs *syd-2* | 0.00001 | Yes |

**Supplementary Table 12: (Associated with Fig. 2J)**

| Analyses of APB-3::GFP puncta - number per cell body, size and intensity | | | |
|---|---|---|---|
| **Analysis** | **Genotype** | **P-value** | **Significant** |
| | **Mann-Whitney Test** | | |
| Average mean puncta intensity to cytosolic intensity per cell body | WT vs *lrk-1* | 0.03486 | Yes |
| | WT vs *syd-2* | 0.00001 | Yes |
| | *lrk-1* vs *syd-2* | 0.00001 | Yes |

**Supplementary Table 13: (Associated with Fig. 3D)**

| Fraction of SNG-1 co-migrating with CTNS-1 | | |
|---|---|---|
| **Genotype** | **P-value** | **Significant** |
| **Mann-Whitney Test** | | |
| WT vs *unc-104(e1265tb120)* | 0.85 | No |
| WT vs *lrk-1; unc-104(e1265tb120)* | 0.015 | Yes |
| WT vs *apb-3; unc-104(e1265tb120)* | 0.015 | Yes |
| *lrk-1(km17)* vs *lrk-1; unc-104(e1265tb120)* | 1.1E-06 | Yes |
| *apb-3(ok429)* vs *apb-3; unc-104(e1265tb120)* | 7.6E-09 | Yes |
| *unc-104* vs *lrk-1; unc-104(e1265tb120)* | 0.005 | Yes |
| *unc-104* vs *apb-3; unc-104(e1265tb120)* | 0.008 | Yes |

**Supplementary Table 14: (Associated with Fig. 4A)**

| Fraction of SNG-1 co-migrating with CTNS-1 | | |
|---|---|---|
| **Genotype** | **P-value** | **Significant** |
| **Mann-Whitney Test** | | |
| WT vs *syd-2(ok217)* | 0.29 | No |
| WT vs *lrk-1; syd-2(ok217)* | 7.4E-08 | Yes |
| WT vs *apb-3; syd-2(ok217)* | 3.6E-01 | Yes |
| WT vs *lrk-1 apb-3; syd-2* | 7.2E-08 | Yes |
| *lrk-1(km17)* vs *lrk-1; syd-2(ok217)* | 0.047 | Yes |
| *lrk-1(km17)* vs *lrk-1 apb-3; syd-2* | 0.11 | No |
| *apb-3* vs *apb-3; syd-2(ok217)* | 4.4E-09 | Yes |
| WT vs *syd-2(ju37)* | 0.39 | No |
| WT vs *apb-3; syd-2(ju37)* | 0.007 | Yes |
| WT vs *lrk-1; syd-2(ju37)* | 5.6E-06 | Yes |
| *syd-2(ju37)* vs *apb-3; syd-2(ju37)* | 0.03 | Yes |
| *syd-2(ju37)* vs *lrk-1; syd-2(ju37)* | 2.5E-05 | Yes |
| *lrk-1(km17)* vs *lrk-1; syd-2(ju37)* | 0.98 | No |
| *apb-3; syd-2(ok217)* vs *apb-3; syd-2(ju37)* | 0.04 | Yes |
| *syd-2(ok217)* vs *syd-2(ju37)* | 0.01 | Yes |
| **One Way ANOVA with Tukey's post Hoc test** | | |
| *apb-3* vs *lrk-1 apb-3* | 0.062 | No |
| *apb-3* vs *lrk-1 apb-3; syd-2* | 0.25 | No |
| *apb-3* vs *apb-3; syd-2(ju37)* | 7.9E-07 | Yes |
| *lrk-1; syd-2(ok217)* vs *lrk-1; syd-2(ju37)* | 0.025 | Yes |
| *lrk-1 apb-3* vs *lrk-1 apb-3; syd-2(ok217)* | 0.4734 | No |

**Supplementary Table 15: (Associated with Fig. 4B)**

| **Fraction of SNG-1 co-migrating with RAB-7** | | |
|---|---|---|
| **Genotype** | **P-value** | **Significant** |
| **Mann-Whitney Test** | | |
| WT vs *apb-3; syd-2* | 0.56 | No |
| *apb-3* vs *apb-3; syd-2* | 0.006 | Yes |
| *syd-2* vs *apb-3; syd-2* | 0.11 | No |
| **One Way ANOVA with Tukey's post Hoc test** | | |
| WT vs *syd-2(ok217)* | 0.88 | No |
| *syd-2* vs *lrk-1* | 0 | Yes |
| *syd-2* vs *apb-3* | 9.1E-08 | Yes |
| WT vs *lrk-1; syd-2* | 6.9E-09 | Yes |
| *lrk-1; syd-2* vs *lrk-1* | 7.3E-01 | Yes |
| *lrk-1; syd-2* vs *syd-2* | 3.8E-07 | Yes |

**Supplementary Table 16: (Associated with Fig. S3C)**

| Percentage vesicles co-migrating CTNS-1 and SNB-1 | | |
|---|---|---|
| **Genotype** | **P-value** | **Significant** |
| **Mann-Whitney test** | | |
| WT vs *syd-2* | 0.39 | No |

**Supplementary Table 17: (Associated with Fig. S3D)**

| Percentage vesicles co-migrating CTNS-1 and RAB-3 | | |
|---|---|---|
| **Genotype** | **P-value** | **Significant** |
| **Mann-Whitney test** | | |
| WT vs *syd-2* | 1 | No |

**Supplementary Table 18: (Associated with Fig. 4G)**

| Percentage vesicles co-migrating SNB-1 and RAB-3 | | |
|---|---|---|
| **Genotype** | **P-value** | **Significant** |
| **One Way ANOVA with Tukey's post Hoc test** | | |
| WT vs *syd-2* | 1.50E-08 | Yes |

**Supplementary Table 19: (Associated with Fig. 4H)**

| Fraction of SNG-1 co-migrating with RAB-3 | | |
|---|---|---|
| **Genotype** | **P-value** | **Significant** |
| **One Way ANOVA with Tukey's post Hoc test** | | |
| WT vs *syd-2*(*ok217*) | 0.49 | No |
| *lrk-1* vs *syd-2* | 1.8E-04 | Yes |
| *apb-3* vs *syd-2* | 0.78 | No |

**Supplementary Table 20: (Associated with Fig. 5D)**

| Fraction of SNG-1 co-migrating with CTNS-1 | | |
|---|---|---|
| **Genotype** | **P-value** | **Significant** |
| **Mann-Whitney Test** | | |
| WT vs *unc-104(e1265tb120); syd-2(ok217)* | 0.95 | No |
| *syd-2(ok217)* vs *unc-104(e1265tb120); syd-2* | 0.28 | No |
| *unc-104(e1265tb120)* vs *unc-104; syd-2(ok217)* | 0.57 | No |

**Supplementary Table 21: (Associated with Fig. 6G)**

| Analyses of UNC-101::GFP puncta - intensity, size and number per cell body | | | |
|---|---|---|---|
| **Analysis** | **Genotype** | **P-value** | **Significant** |
| **One Way ANOVA with Tukey's post Hoc test** | | | |
| Average mean puncta intensity to cytosolic intensity per cell body | WT vs *syd-2(ok217)* | 7.1E-08 | Yes |

**Supplementary Table 22: (Associated with Fig. 6H)**

| Analyses of UNC-101::GFP puncta - intensity, size and number per cell body | | | |
|---|---|---|---|
| **Analysis** | **Genotype** | **P-value** | **Significant** |
| | **Mann-Whitney Test** | | |
| Puncta size average per cell body | WT vs *syd-2(ok217)* | 1.1E-03 | Yes |

**Supplementary Table 23: (Associated with Suppl. Fig. S4D)**

| Analyses of UNC-101::GFP puncta - intensity, size and number per cell body | | | |
|---|---|---|---|
| **Analysis** | **Genotype** | **P-value** | **Significant** |
| | **Mann-Whitney Test** | | |
| Number of puncta per cell body | WT vs *syd-2(ok217)* | 0.07 | No |

**Supplementary Table 24: (Associated with Fig. 6J)**

| Analyses of UNC-101::GFP puncta – intensity per cell body | | | |
|---|---|---|---|
| **Analysis** | **Genotype** | **P-value** | **Significant** |
| | **Mann-Whitney Test** | | |
| Average mean puncta intensity to cytosolic intensity per cell body | WT vs *syd-2(ok217)* | 0.033 | Yes |

**Supplementary Table 25: (Associated with Suppl. Fig. S4F)**

| Analyses of average size of UNC-101::GFP puncta per cell body | | | |
|---|---|---|---|
| **Analysis** | **Genotype** | **P-value** | **Significant** |
| | **Mann-Whitney Test** | | |
| Number of puncta per cell body | WT vs *syd-2(ju37)* | 0.134 | No |

**Supplementary Table 26: (Associated with Suppl. Fig. S4G)**

| Analyses of intensity of UNC-101::GFP puncta per cell body | | | |
|---|---|---|---|
| **Analysis** | **Genotype** | **P-value** | **Significant** |
| | **Mann-Whitney Test** | | |
| Number of puncta per cell body | WT vs *syd-2(ju37)* | 0.97 | No |

Bounoutas, A., Zheng, Q., Nonet, M. L. and Chalfie, M. (2009) 'mec-15 encodes an F-box protein required for touch receptor neuron mechanosensation, synapse formation and development', *Genetics,* 183(2), pp. 607-4SI.
Brenner, S. (1974) 'The genetics of Caenorhabditis elegans', *Genetics,* 77(1), pp. 71-94.
Choudhary, B., Kamak, M., Ratnakaran, N., Kumar, J., Awasthi, A., Li, C., Nguyen, K., Matsumoto, K., Hisamoto, N. and Koushika, S. P. (2017) 'UNC-16/JIP3 regulates early events in synaptic vesicle protein trafficking via LRK-1/LRRK2 and AP complexes', *PLOS Genetics,* 13(11), pp. e1007100.
Consortium, T. C. e. D. M. (2012) 'Large-Scale Screening for Targeted Knockouts in the Caenorhabditis elegans Genome', *G3 Genes|Genomes|Genetics,* 2(11), pp. 1415-1425.
Dwyer, N. D., Adler, C. E., Crump, J. G., L'Etoile, N. D. and Bargmann, C. I. (2001) 'Polarized Dendritic Transport and the AP-1 μ1 Clathrin Adaptor UNC-101 Localize Odorant Receptors to Olfactory Cilia', *Neuron,* 31(2), pp. 277-287.
Kaplan, O. I., Molla-Herman, A., Cevik, S., Ghossoub, R., Kida, K., Kimura, Y., Jenkins, P., Martens, J. R., Setou, M., Benmerah, A. and Blacque, O. E. (2010) 'The AP-1 clathrin adaptor facilitates cilium formation and functions with RAB-8 in C. elegans ciliary membrane transport', *Journal of cell science,* 123(Pt 22), pp. 3966-3977.
Kumar, J., Choudhary, B. C., Metpally, R., Zheng, Q., Nonet, M. L., Ramanathan, S., Klopfenstein, D. R. and Koushika, S. P. (2010) 'The Caenorhabditis elegans Kinesin-3 Motor UNC-104/KIF1A Is Degraded upon Loss of Specific Binding to Cargo', *PLOS Genetics,* 6(11), pp. e1001200.
Lee, J., Jongeward, G. D. and Sternberg, P. W. (1994) 'unc-101, a gene required for many aspects of Caenorhabditis elegans development and behavior, encodes a clathrin-associated protein', *Genes Dev,* 8(1), pp. 60-73.
Maniar, T. A., Kaplan, M., Wang, G. J., Shen, K., Wei, L., Shaw, J. E., Koushika, S. P. and Bargmann, C. I. (2012) 'UNC-33 (CRMP) and ankyrin organize microtubules and localize kinesin to polarize axon-dendrite sorting', *Nature Neuroscience,* 15(1), pp. 48-56.
Nonet, M. L. (1999) 'Visualization of synaptic specializations in live C. elegans with synaptic vesicle protein-GFP fusions', (0165-0270 (Print)).
Sakaguchi-Nakashima, A., Meir, J. Y., Jin, Y., Matsumoto, K. and Hisamoto, N. (2007) 'LRK-1, a C. elegans PARK8-related kinase, regulates axonal-dendritic polarity of SV proteins', *Curr Biol,* 17(7), pp. 592-8.
Zhen, M. and Jin, Y. (1999) 'The liprin protein SYD-2 regulates the differentiation of presynaptic termini in C. elegans', *Nature,* 401(6751), pp. 371-375.
Edwards, S. L., Yu, S. C., Hoover, C. M., Phillips, B. C., Richmond, J. E. and Miller, K. G. (2013) 'An organelle gatekeeper function for Caenorhabditis elegans UNC-16 (JIP3) at the axon initial segment', *Genetics,* 194(1), pp. 143-61.
Zhao, H. and Nonet, M. L. (2001) 'A Conserved Mechanism of Synaptogyrin Localization', *Molecular biology of the cell,* vol. 12 (2001)(8), pp. 2275-89.
